# HieraPTM: From PTM Site Recognition to Functional Phosphosite Prediction through Hierarchical Context and Protein Mechanics

**DOI:** 10.64898/2026.11.12.766127

**Authors:** Yuzhi Yuan, Di Wang, Pengyu Chen, Yanze Niu, Zhiyuan Yang, Hengjia Zhang, Xicheng He, Hao Li, Ziwei Su, Changwei Lin, Tianhang Chen, Chang Ye, Tianchi Lu

## Abstract

Protein post-translational modifications (PTMs) regulate cellular signaling and protein function, making accurate site identification essential for understanding protein regulation. However, PTM site recognition and functional phosphosite prediction are largely modeled independently, limiting the transfer of large-scale recognition knowledge to functional prioritization. Here, we introduce HieraPTM–HieraFuncPhos, a cross-task hierarchical framework that separately models and conditionally integrates transferable PTM recognition representations and function-specific mechanical evidence. HieraPTM integrates local sequence patterns, protein language model representations, and three-dimensional structural information to recognize multiple PTM types. HieraFuncPhos transfers these representations through parameter-efficient adaptation and adaptively retrieves local and distal mechanical evidence conditioned on functional representations. At HSP90AA1 T36 and T115, the model reveals distinct mechanical response patterns consistent with published nuclear magnetic resonance (NMR) observations, suggesting that the framework not only prioritizes functional phosphosites but also provides biophysically grounded, testable hypotheses regarding site-specific regulation. HieraPTM achieved macro-averaged AUROC and AUPRC values of 0.9237 and 0.8425 across eight PTM tasks, while maintaining competitive performance on severely imbalanced external benchmarks. HieraFuncPhos achieved an AUROC of 0.8274 and an AUPRC of 0.8188, outperforming existing comparison methods. Together, this framework connects PTM site recognition, functional prioritization, and structure-informed hypotheses regarding potential regulatory processes.

## 1 Introduction

Protein post-translational modifications (PTMs) participate extensively in biological processes, including cellular signaling and metabolic regulation, by modulating protein activity, stability, conformation, and molecular interactions. Their dysregulation is closely associated with various human diseases[1]. In recent years, rapid advances in mass spectrometry-based proteomics have substantially expanded the identification of diverse PTM sites, including phosphorylation, acetylation, and ubiquitination sites, providing extensive data for systematic studies of protein modifications and their regulation[2]. However, experimental identification of a modification site does not establish its functional importance [3]. The continuing expansion of site discovery, coupled with relatively limited evidence of functional relevance, highlights the need to identify modification sites with potential functional importance [4]. In particular, phosphorylation is an important mechanism of cellular signaling and protein function regulation, supported by extensive experimental data and research. Phosphorylation specificity is influenced by both local substrate sequence motifs and broader molecular recognition mechanisms, including docking interactions [5]. Nevertheless, the functional importance of many identified phosphosites remains to be assessed [6]. Identifying phosphosites with potential functional importance on the basis of multi-type PTM site recognition therefore represents a natural extension from predicting modification occurrence to assessing functional relevance. Although these tasks have different prediction objectives, they may share discriminative information concerning modification acceptors and their sequence and structural contexts. Effectively transferring knowledge learned from large-scale PTM site recognition to functional phosphosite prediction, while incorporating function-specific evidence, is a key challenge in connecting these two prediction tasks.

High-quality training data provide an important foundation for PTM site prediction. DeepMVP [7] established the high-quality PTMAtlas site dataset through standardized reanalysis and quality control of public mass spectrometry datasets and achieved good predictive performance using a CNN–BiGRU sequence model. This study highlighted the importance of data quality and systematic data integration for PTM recognition and provided a reliable foundation for subsequent model development. However, DeepMVP mainly relies on local sequences surrounding candidate residues, and its relatively simple architecture does not explicitly exploit the complementary information contained in protein-wide context and three-dimensional spatial microenvironments. Meanwhile, advances in protein language models and structure prediction have provided richer biological representations for PTM site recognition [8, 9]. For example, PTransIPs incorporates pretrained sequence and structure embeddings into a CNN–Transformer framework for phosphorylation-site prediction, while PDeepPP combines pretrained protein language representations with parallel CNN and Transformer branches for PTM-site and bioactive-peptide classification [10, 11]. By integrating local sequence patterns, protein-wide context, and three-dimensional spatial microenvironments, multimodal models can capture modification-related discriminative information at different levels, offering new opportunities to improve the recognition of complex sites and generalization robustness across different protein backgrounds. However, the complementary value of different information sources does not imply that they contribute equally to all candidate sites. Effectively organizing and integrating this information according to site-specific recognition requirements remains an important challenge in multimodal PTM prediction. Specifically, PTM-CMGMS [12] combines multiscale sequences with structural representations at the residue, atom, and protein levels and enhances feature representations through contrastive learning. However, its multigranularity fusion primarily focuses on the integrated use of structural information without explicitly distinguishing the incremental contribution of structural evidence to existing sequence-based judgments; DeepPTMPred [13] integrates sequence, language model, and structural descriptors for multimodal PTM site prediction. However, it mainly relies on concatenating independently encoded features, without explicitly modeling differences in information requirements among candidate sites or the complementary relationships among multimodal evidence, limiting cross-modal interactions and effective information use; PTM-Mamba [14] integrates PTM-aware representations and protein language model features through bidirectional state-space modeling and dynamic gating. However, its fusion mechanism mainly targets modification-aware protein representation learning and does not specifically characterize local recognition evidence at different candidate sites. Although existing studies have demonstrated the potential of multimodal information for PTM site prediction, biological evidence from different sources has distinct information scales and representational characteristics. How to effectively organize this information around candidate sites and fully exploit complementary advantages across modalities requires further investigation. This motivates more hierarchical integration strategies that fully use biological evidence at different scales and improve the modeling of complex PTM recognition patterns.

Whereas multi-type PTM site recognition primarily focuses on patterns associated with modification occurrence, functional phosphosite prediction assesses the potential functional importance of identified phosphosites. Research on these two tasks has long focused largely on their respective prediction objectives. In early work, Ochoa et al. [15] established functional importance scores for phosphosites by integrating proteomic, structural, regulatory, and evolutionary information. With the development of deep learning, FuncPhos-SEQ [16] used sequence motifs and protein interaction network information for functional prediction, while cDL-FuncPhos [17] and FuncPhos-STR [18] further incorporated protein structural and dynamic features, expanding the biophysical evidence available for functional site identification. More recently, MMFuncPhos [19] enhanced functional representations using protein language models, local structural graphs, and cross-modal attention. PhosLLPS [20] used graph neural networks to identify functional phosphosites involved in regulating liquid–liquid phase separation, extending functional prediction toward multimodal learning and specific biological processes. Despite continued improvements in functional site prediction, multi-type PTM site recognition and functional phosphosite prediction have largely followed relatively independent modeling paths. How the representational knowledge accumulated through large-scale PTM site recognition can support functional importance assessment remains insufficiently explored. Previous studies have indicated that modification occurrence is jointly influenced by acceptor residues, local sequence motifs, and protein structural environments, whereas the functional effects of modifications are also closely related to protein interactions, conformational states, and dynamic regulatory processes [21]. These biological connections and differences motivate knowledge transfer from multi-type PTM site recognition to functional phosphosite prediction, together with the integration of specific evidence related to site functional importance on the basis of shared representations.

Accordingly, we designed a cross-task hierarchical prediction framework comprising HieraPTM and Hiera-FuncPhos. First, using the high-quality PTMAtlas data provided by DeepMVP, we developed HieraPTM to cover eight residue-specific PTM types. Multiscale CNN–BiGRU encoding, position-wise and channel-wise residual gating, and a center-aware Transformer selectively fuse local sequence and protein context. Local three-dimensional residue graphs and structure-guided residual correction further enable hierarchical integration of biological information across scales. We then introduce HieraFuncPhos, which transfers PTM site recognition representations to functional phosphosite prediction. Through LoRA-based parameter-efficient functional adaptation [22] and function-conditioned multiscale mechanical evidence retrieval, it selectively integrates local and distal mechanical information, establishing a cross-task prediction framework that combines shared recognition knowledge with function-specific evidence. This study improves the accuracy of PTM site recognition and functional phosphosite prediction, reveals the distinct contributions of biological evidence at different scales to site recognition, and explores the value of transferring modification recognition knowledge to functional importance assessment, providing a hierarchical computational framework connecting PTM site discovery with functional relevance assessment.

## 2 Materials and Methods

### 2.1 Datasets and Preprocessing

We constructed the PTM site prediction dataset using the human PTMAtlas data released by DeepMVP [7], covering acetylation (lysine, K), methylation (lysine and arginine, K/R), N-glycosylation (asparagine, N), phosphorylation (STY), sumoylation (K), and ubiquitination (K). These were further divided into eight residue-specific prediction tasks according to the acceptor residues. Protein accession numbers, amino acid sequences, and site coordinates were consistently referenced to the UniProtKB/Swiss-Prot release dated August 3, 2022[23]. Candidate site definitions were directly adopted from the dataset released by DeepMVP. Positive samples were PTMAtlas sites supported by MS/MS experimental evidence. Candidate negative samples were constructed from residues of the corresponding acceptor type in the same proteins that lacked known PTM evidence, with known PTM sites already included in the positive dataset or public databases such as PhosphoSitePlus [24] and UniProt further excluded.

For each candidate site, a local sequence window of 61 residues centered on the target residue was extracted, including 30 amino acids on each side, with positions beyond the protein boundaries padded with X. After sequence and site consistency checks, the final dataset comprised 1,116,388 candidate PTM sites from 16,058 human proteins, including 358,022 positive sites and 758,366 candidate negative sites.

To obtain contextualized protein sequence representations, we used ESM-2 to extract per-residue representations from full-length Swiss-Prot protein sequences and cropped the 61 *×* 640 feature matrices corresponding to the 61-residue candidate windows[25]. Proteins exceeding the maximum model input length were processed using overlapping chunks. Valid ESM-2 representations were obtained for 1,116,190 candidate sites, accounting for 99.98% of all samples.

Three-dimensional protein structures were obtained from AlphaFold Protein Structure Database human proteome version 6[26, 27]. Through sequence matching and residue coordinate mapping, candidate PTM sites were located in the corresponding AlphaFold structures. Local three-dimensional residue graphs were constructed around the candidate-site C*α* atom, retaining at most 32 residues from a 12 Å spatial neighborhood as structural input. Valid three-dimensional structural representations were obtained for 976,513 candidate sites, accounting for 87.47% of all samples.

For data partitioning, 10% of samples were reserved in advance as an independent test set and kept isolated throughout model development. The remaining 90% formed the development set for architecture design, hyperparameter optimization, and model selection. After the final model was determined, the development set was further divided at a 9:1 ratio, yielding a final training set comprising 81% of all samples and a validation set comprising 9%. The independent test set remained isolated and was used only for final performance evaluation.

To further examine HieraPTM’s generalization to unseen protein sequences and assess the potential influence of training–test sequence similarity on predictive performance, we constructed an additional protein-disjoint evaluation dataset beyond the original site-level comparison benchmark. The data were repartitioned using proteins as the basic units, ensuring that proteins in the independent test set did not appear in the training or validation set and that all candidate PTM sites from the same protein belonged to the same partition. This avoided potential information leakage caused by different modification sites from the same protein appearing across partitions.

Following protein-disjoint partitioning, CD-HIT-2D [28] was further used to control sequence similarity between the development set and the independent test set. Using training and validation sequences as references, sequence identity thresholds of 90%, 80%, and 70% were applied separately. Test sequences meeting the corresponding similarity criteria against reference sequences, together with their associated sites, were removed to construct progressively stricter homology-controlled test subsets. Using the unfiltered protein-disjoint test set as the baseline, we compared HieraPTM’s AUROC and AUPRC across similarity thresholds to assess prediction stability on unseen proteins and low-similarity sequences.

Ultimately, all 1,116,388 candidate sites had standardized 61-residue local sequence inputs, 99.98% had valid ESM-2 representations, and 87.47% had available AlphaFold three-dimensional structural information. Local sequences, ESM-2 representations, and three-dimensional structures were deterministically aligned by PTM type, protein accession number, and candidate residue coordinates. Samples with missing ESM-2 or structural modalities were retained and handled using the corresponding validity masks. Sample sizes, positive and candidate negative compositions, data partitions, and multimodal coverage for each PTM task are summarized in Supplementary Table S1.

For HieraFuncPhos, we used experimentally observed STY phosphosites from the FuncPhos-STR [18] release. Sites with curated regulatory annotations or PhosphoSitePlus disease-association annotations were labeled as positive, whereas sites lacking these annotations were labeled as negative and treated as unlabeled background. Details of label construction and sampling are provided in the Supplementary Methods.

### 2.2 Model Design

We propose HieraPTM (Hierarchical Context Integration for PTM Site Prediction), which predicts PTM sites by hierarchically integrating local sequence, protein language model, and three-dimensional structural information (Figure 1). For the eight residue-specific PTM types, separate binary classification models are trained with the same architecture but independent parameters.

**Fig. 1.**
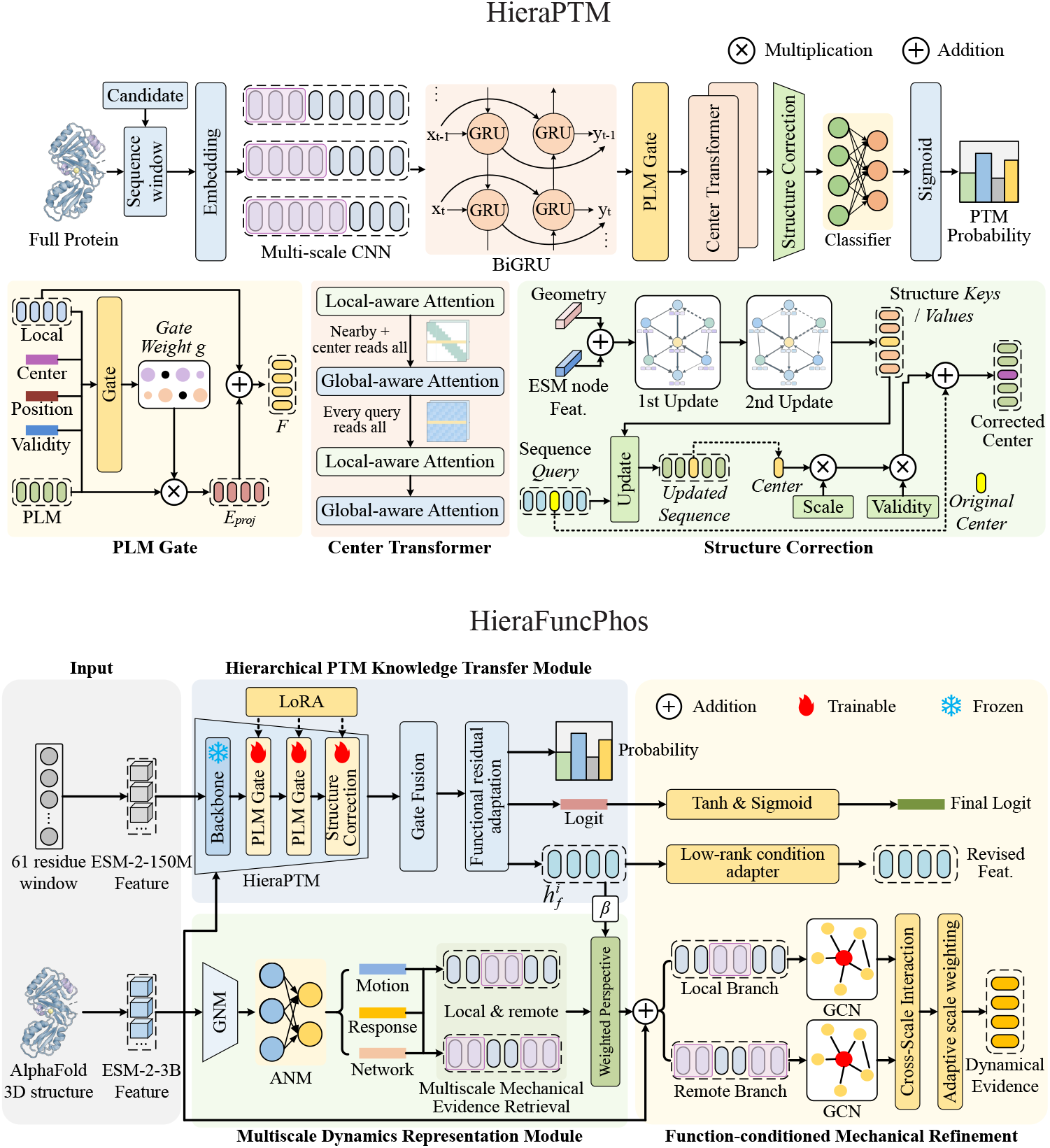
Overview of the HieraPTM and HieraFuncPhos framework. **Top:** HieraPTM hierarchically integrates local sequence patterns, protein language model representations, and three-dimensional structural information for PTM-site recognition. Multiscale CNN–BiGRU extracts local sequence features, while fine-grained ESM-2 gating and a center-aware Transformer incorporate complementary contextual information. Structure-guided cross-attention introduces residual structural corrections to refine site-level predictions. **Bottom:** HieraFuncPhos transfers PTM recognition knowledge through LoRA-based functional adaptation and integrates multiscale mechanical evidence derived from protein structures. Function-conditioned retrieval, local–remote graph modeling, and adaptive cross-scale weighting jointly refine representations for functional phosphosite prediction.

HieraPTM first uses multiscale convolutions and a bidirectional GRU to encode local sequence patterns and bidirectional residue dependencies around candidate sites; it then selectively integrates contextualized representations from frozen ESM-2 through residual gating and uses a center-aware Transformer to model relationships between the candidate site and surrounding residues. For samples with valid structural information, local three-dimensional residue graphs further encode the spatial microenvironment of the candidate site, and cross-modal attention applies residual correction to the sequence–language representations. The final center-site representation predicts the occurrence probability of the corresponding PTM.

#### 2.2.1 Local sequence-context encoding

For each candidate site, the input 61-residue local sequence is represented as

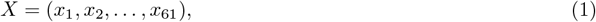

where the 31st residue is the candidate center site, i.e., *c* = 31. Each amino acid is first mapped to a 64-dimensional trainable embedding and then passed to three one-dimensional convolutional branches with kernel sizes of 3, 5, and 7. Each branch has 64 output channels and uses the GELU activation function. Representations from the three scales are concatenated along the channel dimension and normalized using LayerNorm, yielding

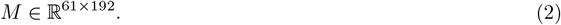

A single-layer bidirectional GRU[29] then further models bidirectional residue dependencies within the local sequence. Each direction contains 64 hidden units. Their outputs are linearly projected to 192 dimensions and fused with the convolutional representation through a residual connection:

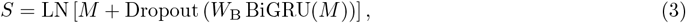

which provides the local sequence representation for subsequent language model fusion. For padding positions introduced at protein termini, a validity mask excludes them from subsequent recurrent encoding, attention computations, and aggregation.

#### 2.2.2 PLM-enhanced center-aware representation

To supplement protein context beyond the fixed local window, HieraPTM uses the 150M-parameter ESM-2 model[25] to extract per-residue representations from full-length proteins and crops the features corresponding to the candidate window. The final ESM-2 layer produces

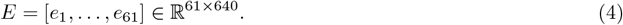

This representation is mapped through a 640 256 192 projection network to the same latent space as the local sequence branch:

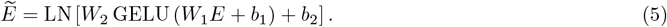

HieraPTM selectively integrates ESM-2 information using a position-wise and channel-wise residual gating mechanism:

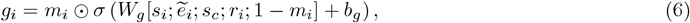

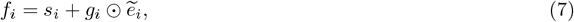

where *s*_*c*_ is the local sequence representation of the center site, *r*_*i*_ is the positional encoding of position *i* relative to the candidate center, and *m*_*i*_ ∈ {0, 1} is the ESM-2 validity mask. The gate bias is initialized to − 2.0, so that the model primarily retains task-specific local sequence representations early in training. Whole-modality ESM-2 dropout with probability 0.1 is additionally applied during training of the sequence–language backbone to reduce excessive reliance on pretrained representations.

The fused representations are further passed to a four-layer center-aware Transformer[30]. The encoder has a hidden dimension of 192, 6 attention heads, and a 512-dimensional feed-forward network, with an alternating local–global–local–global attention pattern. In local attention layers, ordinary positions interact only with valid residues within a sequence distance of 8, whereas the candidate center can access the entire valid window. To formalize this asymmetric attention mechanism, the access mask for local attention layers is defined as

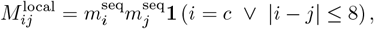

where 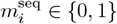 indicates the validity of a sequence position.

When the query corresponds to the candidate center, i.e., *i* = *c*, it can access all valid keys; ordinary queries can access only valid residues within a sequence distance of 8. This special access is directional: distant ordinary queries do not automatically gain access to the center.

Correspondingly, the mask for global attention layers is

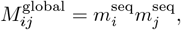

allowing attention interactions among all valid residues. The model also introduces learnable absolute positional embeddings and binary center-state embeddings to distinguish the candidate PTM center residue from ordinary sequence positions.

To explicitly describe the positional relationships of residues relative to the candidate center, signed relative positional biases and center-relation biases are added to standard scaled dot-product attention:

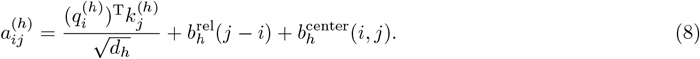

Here, the signed relative positional bias is defined according to the position of the key relative to the query, *d*_*ij*_ = *j − i*. For a sequence window of length *L* = 61, relative distances range over [*−*60, 60], corresponding to 2*L −* 1 = 121 learnable position buckets:

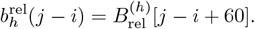

The center-relation bias further distinguishes four relationships between the query and key relative to the candidate center. The relation index is defined as

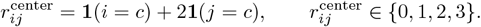

where the four states correspond, respectively, to noncenter–noncenter, center query–noncenter key, noncenter query–center key, and center–center relationships.

Each attention head has an independent learnable four-state bias:

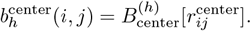

These two types of bias jointly modify attention scores, which are normalized by softmax under the constraints of the local or global access mask.

After four encoding layers, we obtain

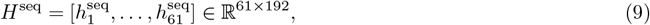

and extract the representation corresponding to the candidate center,

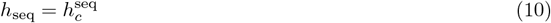

as the site-level representation of the sequence–language branch.

#### 2.2.3 Local 3D microenvironment encoding

For candidate sites with valid structural information, HieraPTM further constructs a local three-dimensional residue graph centered on the candidate residue,

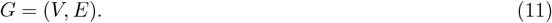

retaining at most 32 neighboring residues within a 12 Å spatial neighborhood of the candidate-center C*α* atom, which form the local graph together with the center residue.

Each node contains both a frozen ESM-2 residue representation and local geometric descriptors. ESM-2 node features are mapped to 192 dimensions using the same projection network as the sequence branch, while geo-metric features are nonlinearly projected to the same latent space. The two are added to obtain the initial node representation:

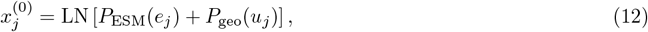

where *u*_*j*_ denotes the local geometric features of node *j*, including structural confidence, spatial and sequence relationships to the candidate center, and neighborhood information at different distance scales.

The local graph is encoded using two four-head graph attention layers[31]. A message-passing connection is established if two nodes have a C*α* distance no greater than 8 Å, are adjacent in the primary sequence, or are the same node. Edge features are defined as

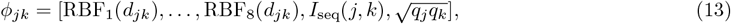

where *d*_*jk*_ is the inter-node C*α* distance, *I*_seq_ indicates adjacency in the primary sequence, and *q*_*j*_ and *q*_*k*_ are normalized pLDDT values. Spatial distances are continuously encoded using 8 radial basis functions.

The weights of graph attention head *h* are expressed as

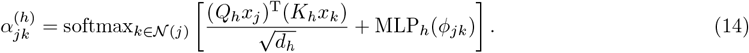

Each graph attention layer incorporates residual connections, LayerNorm, and a feed-forward network. This yields

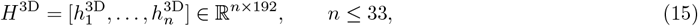

as the local three-dimensional microenvironment representation of the candidate site.

#### 2.2.4 Structure-guided residual correction and PTM prediction

HieraPTM integrates local three-dimensional structural information into the sequence–language representation through cross-modal attention. Sequence tokens serve as queries, while local structural nodes serve as keys and values:

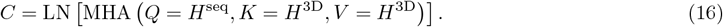

Cross-modal attention contains 4 attention heads. Because the final prediction targets the candidate center residue, only the cross-modal representation *C*_*c*_ corresponding to the center position is retained, and the center-site representation is updated through a residual connection:

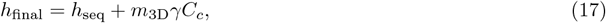

where *m*_3D_ ∈ {0, 1} is the structural validity mask, and *γ* is a learnable structural residual scaling parameter initialized to 0. This initialization preserves the original sequence–language predictions when the structural module is introduced, allowing it to gradually learn incremental corrections from structural information during subsequent training. For samples lacking valid structural information, *m*_3D_ = 0, and the model automatically falls back to the sequence–language pathway.

The final center-site representation is passed through a linear classifier to obtain the prediction logit:

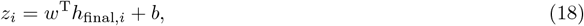

which is converted into the predicted probability of the corresponding PTM using the sigmoid function:

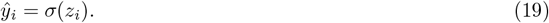

#### 2.2.5 Learning objective and staged optimization

For a training batch containing *N* candidate sites, the true label and predicted probability of sample *i* are denoted by *y*_*i*_ *∈ {*0, 1*}* and *ŷ*_*i*_, respectively. The model is optimized using binary cross-entropy with a positive-class weight:

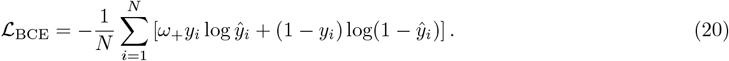

where *ω*_+_ is the positive-class weight used during training for the current PTM task.

### 2.3 Model training and evaluation

HieraPTM uses a two-stage training strategy. The first stage trains the multiscale convolutions, BiGRU, ESM-2 residual gates, center-aware Transformer, and classifier, while pretrained ESM-2 remains frozen throughout; the second stage freezes the sequence–language backbone and classifier and optimizes only the three-dimensional graph encoder, cross-modal attention, and structural residual parameter *γ* to learn incremental corrections to the base predictions from structural information.

During model development, five-fold stratified group cross-validation is used, with a standardized training and evaluation protocol across folds. Early stopping and best-checkpoint selection control training. After the final configuration is determined, the model is retrained and evaluated only on the independent test set reserved in advance. All threshold-dependent metrics use a classification threshold of 0.5.

Model performance is comprehensively evaluated using AUROC, AUPRC, Accuracy, Precision, Recall, Specificity, F1-score, and Matthews correlation coefficient (MCC).

Let TP, TN, FP, and FN denote true positive, true negative, false positive, and false negative, respectively. The metrics are defined as follows:

- **Accuracy:**

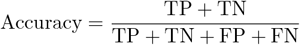
- **Precision:**

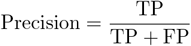
- **Recall (Sensitivity):**

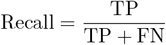
- **Specificity:**

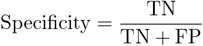
- **F1-score:**

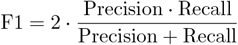
- **Balanced Accuracy (BACC):**

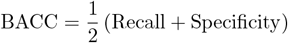
- **Matthews Correlation Coefficient (MCC):**

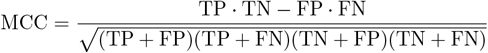
- **AUROC and AUPRC:** the areas under the receiver operating characteristic curve and the precision–recall curve, respectively.

AUROC and AUPRC are calculated from continuous predicted probabilities to assess overall discrimination across decision thresholds; the remaining metrics are calculated from binary predictions at a threshold of 0.5. All metrics are computed independently for each of the eight PTM tasks, and unweighted macro-averages are also reported to give each PTM type equal weight in the overall evaluation. Five-fold cross-validation results are summarized as the mean and standard deviation across folds, whereas final model performance is reported on the independent test set. Further details of the HieraFuncPhos model architecture, training procedures, and multiscale mechanical evidence modeling are provided in the Supplementary Methods.

## 3 Results

### 3.1 Comparative Study

#### 3.1.1 Comparative evaluation on the DeepMVP benchmark

To fairly evaluate the proposed model against existing PTM site prediction methods, we established a standardized site-level comparison benchmark using the DeepMVP dataset, with training, validation, and fixed held-out test sets partitioned at a ratio of 81%/9%/10%. We selected eight representative PTM site prediction methods as baselines: DeepMVP [7], Sitetack [32], PhosIDN [33], DeepPTMPred [13], COMPASS-PTM [34], PTM-Mamba [14], PTM-CMGMS [12], and DeepGpgs [35]. All methods were evaluated on eight PTM prediction tasks except PhosIDN, which was included only in the phosphorylation-ST and phosphorylation-Y comparisons. For each PTM type, all methods used identical data partitions, with the same test samples, labels, and natural class proportions across models. All trainable baseline models were implemented using the authors’ publicly available code or adapted implementations verified for architectural consistency, retrained on the shared training data, and followed their original model designs and training strategies. Model selection, hyperparameter optimization, and decision-threshold determination were restricted to the training and validation sets; the test set was used only for final performance evaluation.

We used AUROC and AUPRC as the primary evaluation metrics and additionally reported Accuracy, Precision, Recall, Specificity, F1, and MCC to comprehensively assess prediction performance across PTM types, particularly classification performance under class imbalance. We also estimated 95% confidence intervals for each model using 2,000 bootstrap resamples to assess the statistical reliability of predictive performance. All metrics were computed consistently on the same fixed test samples.

HieraPTM achieved the highest AUROC on all eight PTM tasks, demonstrating a consistent advantage in discrimination (Figure 2A). Its AUROC reached 0.9969 for N-linked glycosylation-N, indicating effective identification of modification sites with strong sequence characteristics. For acetylation-K, phosphorylation-ST, and phosphorylation-Y, HieraPTM achieved AUROCs of 0.9356, 0.9459, and 0.9096, respectively, outperforming the other models. Compared with the original DeepMVP, HieraPTM increased AUROC for sumoylation-K from 0.8750 to 0.9065, a relative improvement of 3.60%; AUROC increased by 0.0279 and 0.0252 for phosphorylation-Y and methylation-K, respectively. Notably, HieraPTM achieved the highest AUROC even for N-linked glycosylation-N, where predictive performance was already close to saturation. These results indicate that the gains were not restricted to a particular modification type but followed a relatively consistent trend across PTM tasks.

**Fig. 2.**
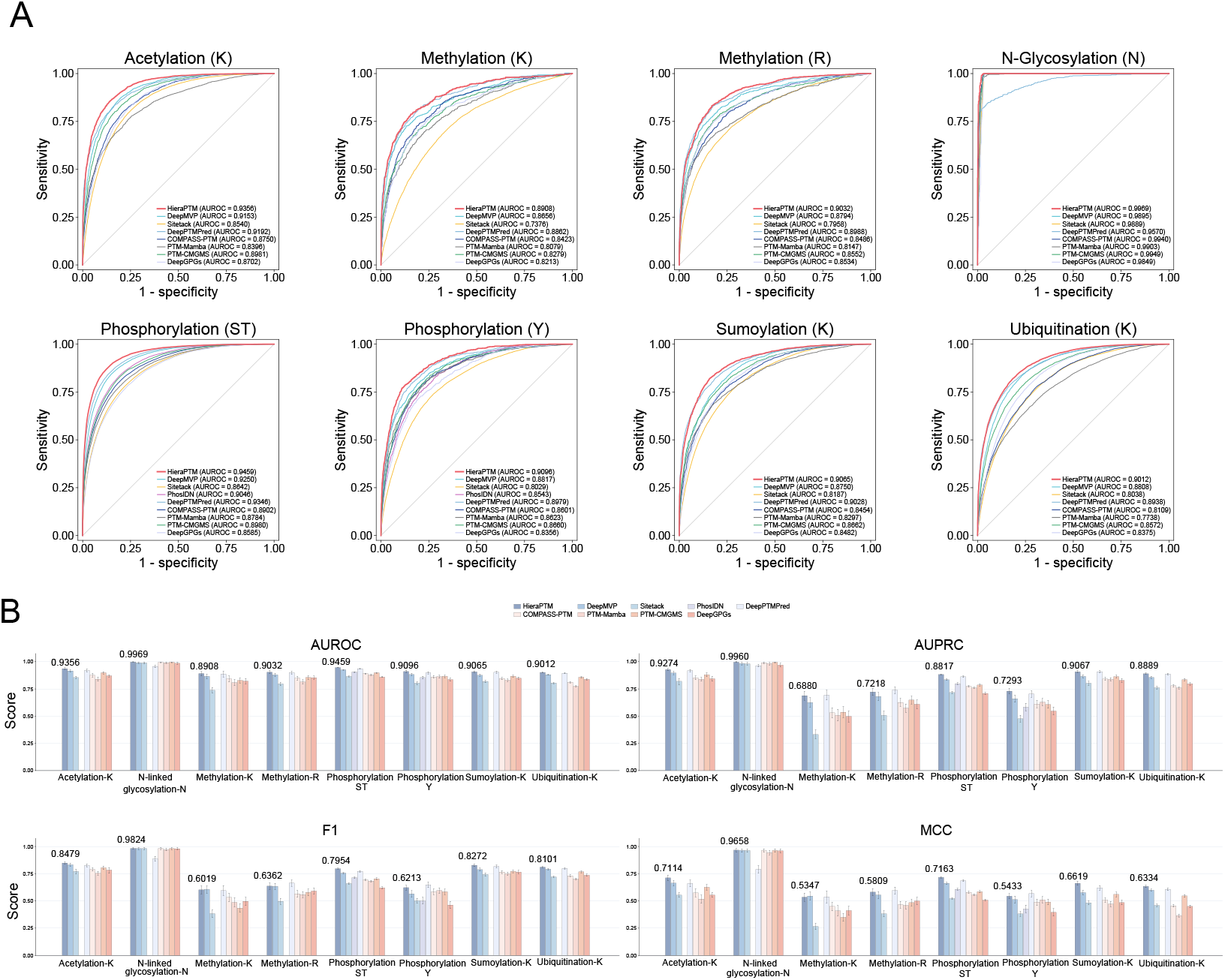
Performance comparison of HieraPTM with existing PTM site predictors on the DeepMVP benchmark. a, Receiver operating characteristic (ROC) curves of HieraPTM and competing methods across eight PTM prediction tasks. b, Task-specific comparisons of AUROC, AUPRC, F1-score and Matthews correlation coefficient (MCC). Error bars indicate 95% confidence intervals estimated from 2,000 bootstrap resamples. PhosIDN was evaluated only on phosphorylation-ST and phosphorylation-Y.

Macro-averaged comparisons further showed that HieraPTM achieved the highest AUROC (0.9237) and AUPRC (0.8425), outperforming all other eight-task prediction models (Table 1). The closest-performing method, DeepPTMPred, achieved 0.9113 and 0.8339, respectively, corresponding to relative AUROC and AUPRC improvements of 1.36% and 1.03% for HieraPTM. Compared with the original DeepMVP, HieraPTM increased AUROC from 0.9015 to 0.9237, a relative improvement of 2.46%, and AUPRC from 0.7997 to 0.8425, a relative improvement of 5.35%. Given the substantial differences in class distributions across PTM tasks, the simultaneous gains in AUROC and AUPRC indicate improvements in both overall discrimination and the ranking of positive modification sites.

**Table 1.** Macro-averaged performance comparison of HieraPTM and existing PTM site predictors on the DeepMVP benchmark. The best result for each metric is highlighted in bold.

| Model | AUROC | AUPRC | Accuracy<br>(%) | F1<br>(%) | Precision<br>(%) | Recall<br>(%) | Specificity<br>(%) | MCC |
|---|---|---|---|---|---|---|---|---|
| DeepMVP | 0.9015 | 0.7997 | 86.19 | 74.37 | **74.84** | 74.85 | 87.63 | 0.6345 |
| Sitetack | 0.8332 | 0.6740 | 79.43 | 65.69 | 61.15 | 72.33 | 80.28 | 0.5011 |
| PhosIDN<sup>*a*</sup> | 0.8794 | 0.6899 | 84.76 | 60.71 | 66.02 | 57.02 | **91.73** | 0.5162 |
| DeepPTMPred | 0.9113 | 0.8339 | 85.63 | 75.06 | 74.07 | 76.49 | 86.75 | 0.6326 |
| COMPASS-PTM | 0.8708 | 0.7503 | 82.34 | 70.51 | 66.08 | 75.87 | 81.52 | 0.5604 |
| PTM-Mamba | 0.8496 | 0.7357 | 81.21 | 68.72 | 65.87 | 72.44 | 80.47 | 0.5290 |
| PTM-CMGMS | 0.8829 | 0.7684 | 81.19 | 70.31 | 64.39 | **80.87** | 81.11 | 0.5754 |
| DeepGPGs | 0.8637 | 0.7249 | 81.17 | 67.93 | 65.27 | 73.86 | 76.86 | 0.5329 |
| **HieraPTM (Ours)** | **0.9237** | **0.8425** | **86.81** | **76.53** | 74.32 | 80.68 | 88.00 | **0.6685** |

*Note:* All models were evaluated across eight PTM prediction tasks, except PhosIDN<sup>*a*</sup>, which covers only phosphorylation-ST and phosphorylation-Y. Its macro-averaged results are therefore not directly comparable with the eight-task averages.

**Table 2.** Macro-averaged performance comparison of HieraPTM and existing PTM site predictors on the dbPTM external-source benchmark across four PTM prediction tasks. The best result for each metric is highlighted in bold.

| Model | AUROC | AUPRC | Accuracy<br>(%) | F1<br>(%) | Precision<br>(%) | Recall<br>(%) | Specificity<br>(%) | BACC<br>(%) | MCC |
|---|---|---|---|---|---|---|---|---|---|
| DeepMVP | 0.8523 | 0.7251 | 74.48 | 68.65 | 58.63 | **88.71** | 66.73 | 77.72 | 0.5302 |
| Sitetack | 0.8184 | 0.6850 | 80.41 | 68.54 | 64.76 | 73.75 | 80.40 | 77.08 | 0.5259 |
| DeepPTMPred | 0.7086 | 0.4716 | 65.65 | 46.76 | 47.63 | 52.88 | 70.83 | 61.86 | 0.2398 |
| COMPASS-PTM | 0.8490 | 0.7236 | 81.58 | 71.86 | 65.80 | 79.72 | 78.28 | 79.00 | 0.5601 |
| **HieraPTM (Ours)** | **0.8692** | **0.7382** | **82.93** | **73.73** | **68.63** | 82.01 | **80.44** | **81.22** | **0.6071** |

**Table 3.** Macro-averaged performance comparison of HieraPTM and existing phosphorylation site predictors on the PhosF3C external-source benchmark across two phosphorylation prediction tasks (Pho-ST and Pho-Y). The best result for each metric is highlighted in bold.

| Model | AUROC | AUPRC | Accuracy<br>(%) | F1<br>(%) | Precision<br>(%) | Recall<br>(%) | Specificity<br>(%) | BACC<br>(%) | MCC |
|---|---|---|---|---|---|---|---|---|---|
| DeepMVP | 0.7081 | 0.2506 | 77.88 | 30.92 | 24.32 | 44.99 | 82.17 | 63.58 | 0.2101 |
| PhosIDN | 0.7143 | 0.2866 | 84.70 | 29.87 | **30.87** | 32.04 | 91.54 | 61.79 | 0.2246 |
| Sitetack | 0.7019 | 0.2611 | 78.81 | 30.49 | 24.17 | 41.86 | 83.61 | 62.74 | 0.2014 |
| DeepPTMPred | 0.6404 | 0.1657 | 74.96 | 21.37 | 20.02 | 32.16 | 80.56 | 56.36 | 0.1096 |
| COMPASS-PTM | 0.7003 | 0.2678 | 81.55 | 31.22 | 26.90 | 37.59 | 87.24 | 62.41 | 0.2139 |
| DeepGPGs | 0.6864 | 0.2430 | **84.77** | 26.53 | 28.96 | 25.88 | **92.39** | 59.13 | 0.1872 |
| **HieraPTM (Ours)** | **0.7374** | **0.3338** | 78.31 | **35.11** | 26.53 | **52.03** | 81.73 | **66.88** | **0.2565** |

HieraPTM also demonstrated strong overall performance on threshold-dependent classification metrics. Its macro-averaged Accuracy, F1-score, and MCC reached 86.81%, 76.53%, and 0.6685, respectively, the highest among the eight-task prediction models (Table 1). Compared with DeepPTMPred, HieraPTM improved Accuracy and F1-score by 1.18 and 1.47 percentage points, respectively, and MCC by an absolute 0.0358, demonstrating advantages in overall classification accuracy and joint discrimination of positive and negative samples. Compared with the original DeepMVP, HieraPTM increased Recall from 74.85% to 80.68%, a gain of 5.83 percentage points, while Precision was only 0.52 percentage points lower. Meanwhile, F1-score increased by 2.16 percentage points, Specificity by 0.37 percentage points, and MCC by 0.0340. These results indicate that HieraPTM effectively improves the detection of positive PTM sites while largely maintaining precision and negative-site recognition, achieving better overall classification performance. Although PTM-CMGMS had slightly higher Recall than HieraPTM (80.87% versus 80.68%), HieraPTM achieved an F1-score higher by 6.22 percentage points and an MCC higher by 0.0931. This indicates that HieraPTM achieves a better balance between positive-site detection and prediction precision rather than merely maximizing recall. Together, these results demonstrate more balanced and reliable overall predictive performance across evaluation dimensions.

To further assess the potential influence of sequence homology on predictive performance, we examined HieraPTM’s robustness on test sets filtered at similarity thresholds of 90%, 80%, and 70% (Supplementary Fig. S1). As the sequence similarity threshold decreased, the ROC and PR curves for the eight PTM types maintained broadly similar discrimination patterns. These results indicate that HieraPTM retains relatively stable predictive ability when training–test sequence similarity is reduced, further supporting its generalization robustness on low-similarity sequences.

To further validate HieraPTM’s cross-protein generalization and prediction stability, we applied CD-HIT [28] with sequence identity thresholds of 90%, 80%, and 70% on top of the strictest fully protein-disjoint criterion, constructing progressively stricter homology-controlled test sets (Supplementary Fig. S1). As the sequence similarity threshold decreased, ROC and PR curves remained highly consistent across all eight PTM tasks, with only small fluctuations in AUROC and AUPRC. This indicates that HieraPTM maintains good discrimination on unseen proteins and low-homology sequences, and that its learned PTM recognition features support cross-protein transfer and robust generalization.

#### 3.1.2 Out-of-distribution generalization across PTM datasets

To assess generalization across data sources, we constructed external test sets using the official dbPTM [36] benchmark and the publicly released independent test data from PhosF3C [37]. To avoid potential information leakage from training-data overlap, we performed strict deduplication, removing sequences overlapping our training and validation sets and sequences present in both the positive and negative sets (i.e., internal conflicts). We further constructed CD-HIT 70% homology-controlled test subsets by removing test proteins with sequence identity reaching or exceeding 70%, together with all their candidate sites. The resulting datasets covered six PTM tasks: N-linked glycosylation-N, methylation-K, methylation-R, ubiquitination-K, phosphorylation-ST, and phosphorylation-Y, with substantial class imbalance (Supplementary Tables S3, S4). We systematically compared HieraPTM with DeepMVP, Sitetack, DeepPTMPred, and COMPASS-PTM, additionally including PhosIDN and DeepGPGs in the phosphorylation tasks. All models generated predictions without using external test labels for training or adjustment.

On the dbPTM external test set, HieraPTM achieved the highest macro-averaged AUROC (0.8692) and AUPRC (0.7382) across four PTM tasks, exceeding DeepMVP’s 0.8523 and 0.7251, respectively (Figure 3A). HieraPTM also achieved macro-averaged F1-score and MCC values of 0.7373 and 0.6071, respectively, outperforming the other models and indicating good overall discrimination across PTM types and class distributions. After applying the CD-HIT 70% homology restriction, HieraPTM achieved macro-averaged AUROC and AUPRC values of 0.8682 and 0.7332, respectively, with F1-score and MCC values of 0.7330 and 0.6041 (Figure 3B). Compared with the standard external evaluation, AUROC and AUPRC decreased by only 0.0010 and 0.0050, respectively, indicating largely stable overall performance.

**Fig. 3.**
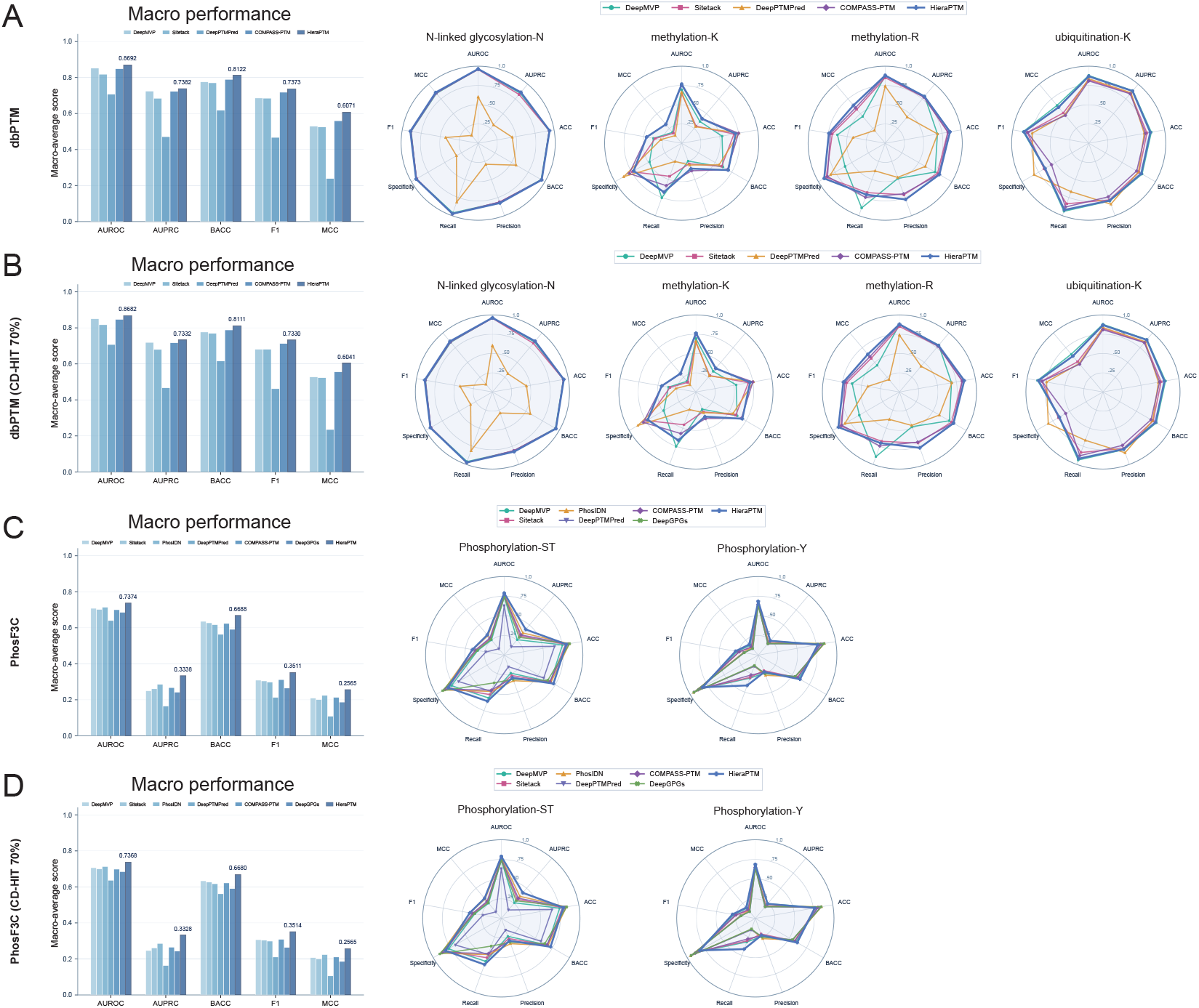
External generalization of HieraPTM across PTM benchmarks. a, Performance comparison on the dbPTM bench-mark across four PTM tasks. b, Homology-controlled evaluation on dbPTM after CD-HIT 70% filtering. c, Performance comparison on the PhosF3C-derived benchmark across phosphorylation-ST and phosphorylation-Y. d, Homology-controlled evaluation on PhosF3C after CD-HIT 70% filtering. Bar charts show macro-averaged performance, while radar plots summarize individual PTM tasks across nine evaluation metrics. All models were evaluated without external test-set fine-tuning, using a fixed threshold of 0.5 for threshold-dependent metrics.

HieraPTM also demonstrated good generalization on the more severely imbalanced PhosF3C external test set. Its macro-averaged AUROC and AUPRC reached 0.7374 and 0.3338, respectively, outperforming the next-best model, PhosIDN (0.7143 and 0.2866), by 0.0231 and 0.0472 (Figure 3C). Given that positive sites accounted for approximately 11% of this dataset, the AUPRC improvement is particularly noteworthy. HieraPTM additionally achieved the highest macro-averaged F1-score (0.3511) and MCC (0.2565), indicating competitive classification performance even at a low positive-site proportion. After CD-HIT 70% filtering, macro-averaged AUROC and AUPRC were 0.7368 and 0.3328, respectively, while F1-score and MCC remained at 0.3514 and 0.2565 (Figure 3D), demonstrating only small performance fluctuations.

### 3.2 Stagewise Separation of PTM-Site Representations

To reveal how integration across information levels affects the organization of PTM site representations, we visualized the feature spaces of all eight PTM types at four successive model stages using UMAP [38] and quantitatively assessed class separability in the original 192-dimensional representations using the Balanced Silhouette Score (Figure 4). As local sequence encoding, protein language model enhancement, center-aware contextual integration, and three-dimensional structural information were successively introduced, PTM site representations exhibited clear stage-dependent evolution.

**Fig. 4.**
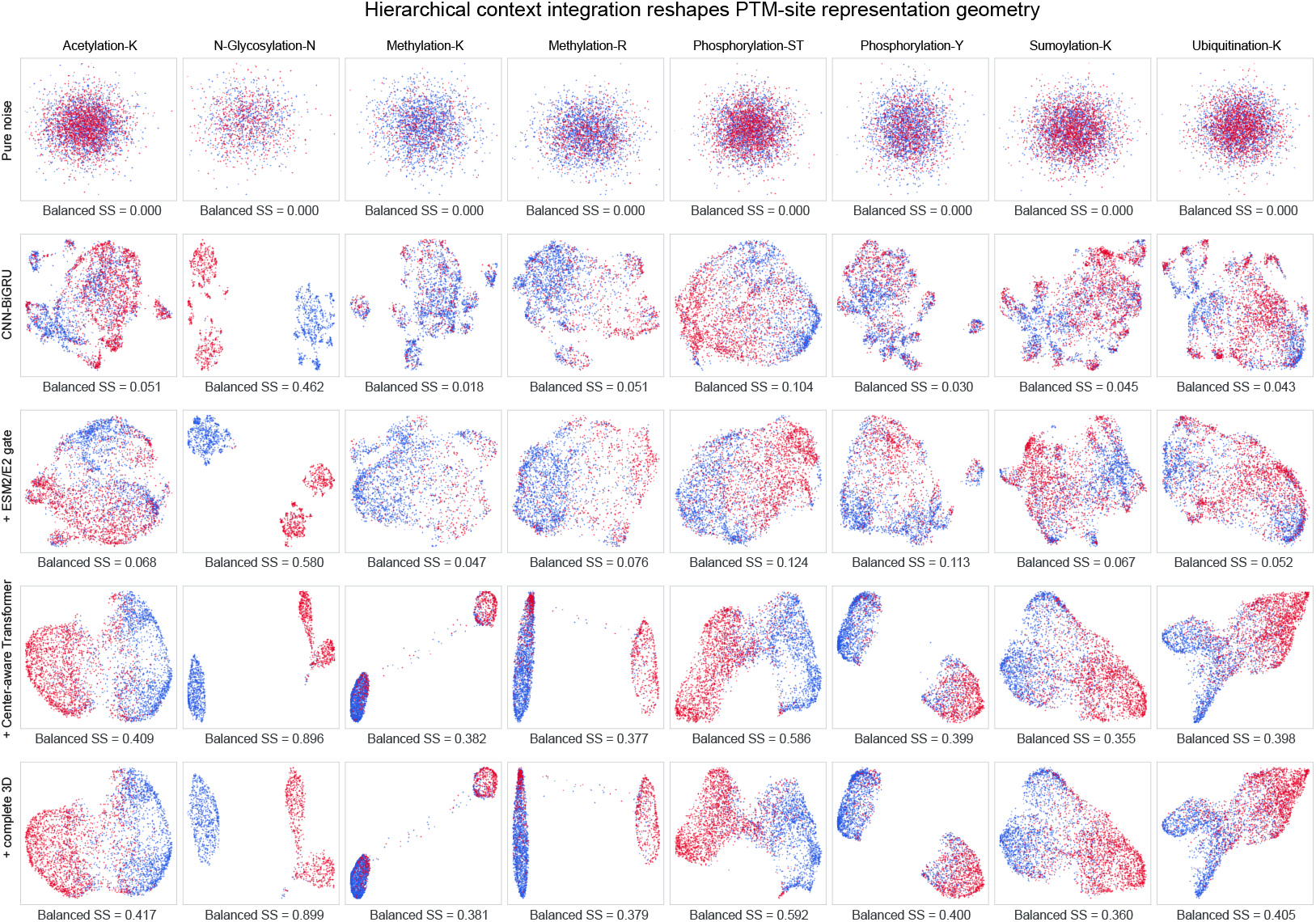
Representation Geometry Evolves across HieraPTM Stages. UMAP visualization of eight PTM types across successive representation stages, from CNN–BiGRU and ESM2 enhancement to Center-aware Transformer and complete 3D integration, with a random-feature control. Red and blue points represent positive and negative PTM sites, respectively.

In the initial CNN–BiGRU representations, positive and negative sites overlapped extensively, indicating that local sequence features had not yet formed well-separated class structures. Introducing ESM2 increased Macro-SS from 0.101 to 0.141, improving class separability while retaining substantial distributional heterogeneity. This reflects the role of protein language models in enriching contextual information while preserving sequence-feature diversity. In contrast, the center-aware Transformer induced more substantial reorganization of the representation space, markedly increasing Macro-SS to 0.475 and producing increasingly compact and distinct discriminative distributions for positive and negative sites. This change indicates that center-aware contextual integration not only strengthens feature associations but also organizes and concentrates distributed contextual information along class-discriminative directions, exhibiting pronounced class-aligned representation compression.

Incorporating complete three-dimensional structural information further increased Macro-SS to 0.479. Unlike the large-scale reorganization at the Transformer stage, integration of three-dimensional information mainly produced subtle adjustments within the existing discriminative space. This difference indicates that structural information complements highly organized sequence representations by providing local spatial environmental information.

### 3.3 Hierarchical context integration reveals distinct mechanisms of PTM recognition

Stage-wise mechanistic analysis showed that the modules perform complementary functions at different scales rather than repeatedly encoding the same information. Center-aware integration first reorganizes local sequence evidence: rather than simply amplifying the importance of residues near the candidate site, it expands contextual utilization while selectively enhancing discriminative residue–position combinations. The mean effect of local grammar disruption increased from 2.609 to 4.877 (Figure 5A), with clear enhancement in 6 of the 8 PTM types. Functional grammars differed substantially across PTMs. The strong local dependence of N-glycosylation was consistent with the classical N-X-S/T sequon recognition mechanism [39]; the importance of acidic residues down-stream of the candidate lysine in sumoylation-K was consistent with UBC9 recognition of *?*KxD/E-like consensus sequences [40]; and the proximal Pro preference observed in Pho-ST agreed with classical recognition patterns of proline-directed Ser/Thr phosphorylation [41]. To examine whether model predictions were consistent with experimentally observed PTM sequence characteristics, we compared neighboring amino acid preferences between experimentally annotated positive sites and HieraPTM-predicted positive sites across the eight PTM types (Supplementary Fig. S3). Sequence logo analysis showed that, after excluding the center residue, the two groups retained highly consistent neighboring amino acid preferences across PTM tasks and preserved characteristic sequence patterns of multiple PTM types. Thus, HieraPTM predictions were highly consistent with the sequence backgrounds of real PTM sites, supporting the biological plausibility of the learned modification recognition patterns. Further bidirectional counterfactual interventions showed that disrupting positive-compatible grammar reduced correct-class evidence in all 8 PTM types, whereas restoring the same grammar produced changes in the opposite direction (Figure 5B). This indicates that the model learned a set of PTM-specific functional recognition grammars with different strengths of local constraints rather than a single shared motif.

**Fig. 5.**
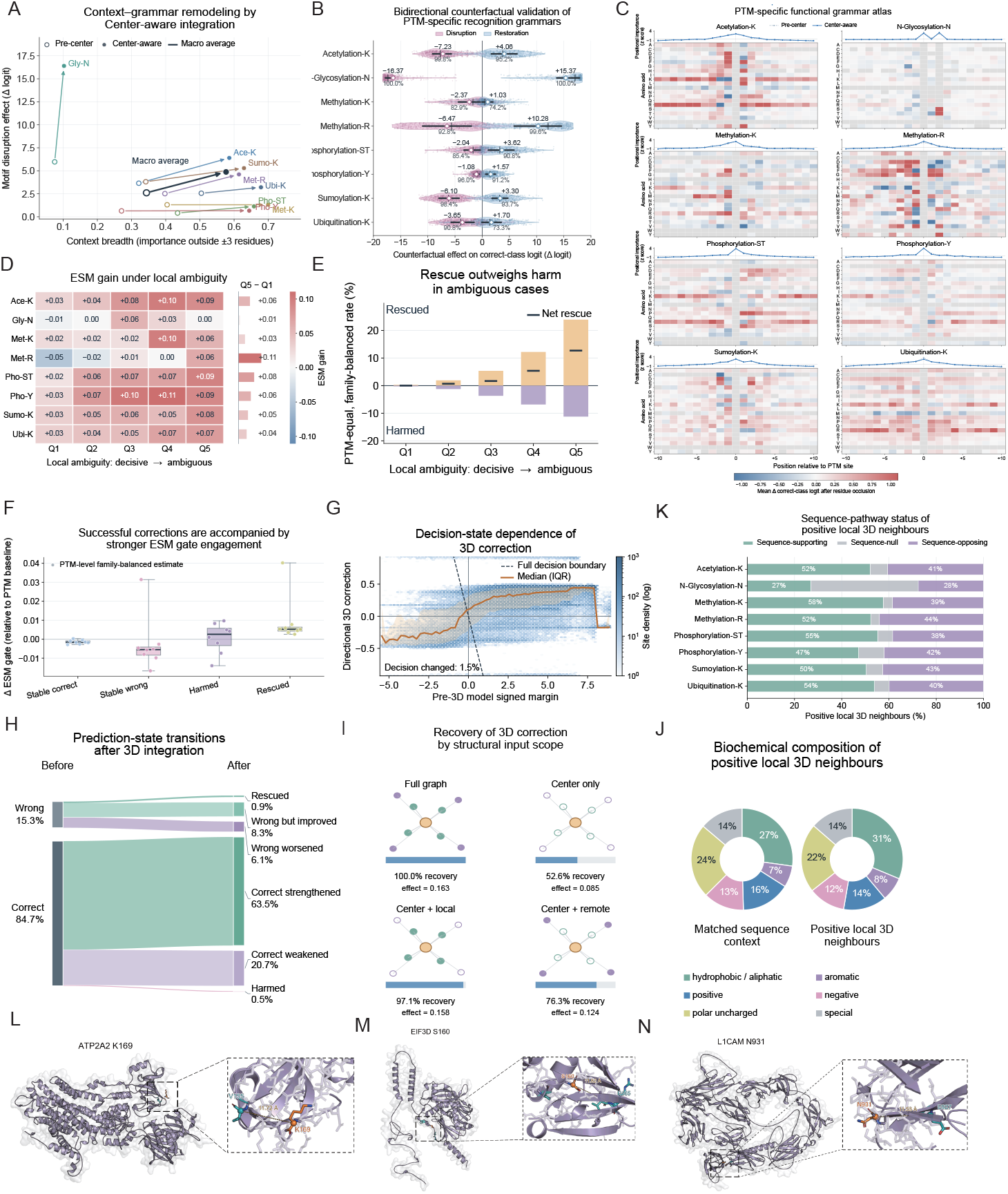
Hierarchical context integration reveals complementary mechanisms of PTM site recognition. a, Context–grammar remodeling from the pre-center to Center-aware representation. b, Bidirectional counterfactual effects of grammar disruption and restoration across eight PTM types. c, PTM-specific residue–position functional grammar atlas. d, ESM gain across local-ambiguity quintiles (Q1, most decisive; Q5, most ambiguous). e, Rescued and harmed prediction transitions across ambiguity levels. f, ESM-gate engagement across prediction outcomes. g, Directional 3D correction across the pre-3D decision margin. h, Prediction-state transitions after 3D integration. i, Recovery of the full 3D correction using different structural input scopes. j, Biochemical composition of positive local 3D neighbours. k, Their relationship to sequence-pathway evidence. l–n, Representative structural examples of ATP2A2 K169, EIF3D S160 and L1CAM N931. Mechanistic analyses were performed on frozen models using independent test data.

When local grammar was insufficient to support stable decisions, higher-level protein context began to compensate. As samples progressed from the most certain Q1 to the most ambiguous Q5, the family-balanced log-loss gain from ESM representations increased from 0.013 to 0.068, with positive Q5–Q1 differences in all 8 PTM types. In Q5, 23.9% of predictions were corrected and 11.2% were adversely affected, yielding a net correction of 12.7 percentage points; successfully corrected classes also exhibited higher ESM gate engagement (Figure 5G). Thus, ESM2 acts as a conditional protein-context compensator: its marginal contribution is limited when local sequence grammar is already clear, whereas protein-level semantic context provides additional discriminative information when local evidence is weak or conflicting.

Three-dimensional structure provides further, more localized residual corrections. Complete 3D fusion changed only approximately 1.48% of binary decisions, mainly affecting samples near the pre-3D decision boundary. When structural inputs were restricted, center-only input recovered only approximately half of the full-graph correction, whereas adding local spatial neighbors recovered approximately 97.1% (Figure 5H, I). This indicates that effective structural information mainly originates from the local spatial microenvironment surrounding the candidate site. The primary role of the 3D branch is to reweight local residues according to spatial proximity in the folded structure.

This spatial compensation mechanism was supported by external biological evidence at representative sites (Figure 5L-N). For ATP2A2 K169, the structural branch can supplement weak sequence evidence using the local spatial neighborhood. Previous studies have shown that JOSD2 can regulate SERCA2a K169 ubiquitination, affecting its stability and cardiomyocyte Ca^2+^ handling and providing independent support for the functional relevance of this site [42]. L1CAM N931 has more direct experimental site-level evidence: site-specific N-sialoglycoproteomics explicitly identified this site and showed that it lies within an N-glycosylation consensus motif [43]. External evidence for EIF3D S160 mainly comes from the human HCC phosphoproteomics dataset PXD028593, providing independent proteomic evidence of phosphorylation at this site [44].

Together, HieraPTM supports a hierarchical PTM recognition mechanism: center-aware modeling first organizes PTM-specific local functional grammars; ESM2 provides protein-context compensation when local rules are insufficient; and the local three-dimensional neighborhood further corrects boundary samples through spatial selection and residue reweighting.

### 3.4 Ablation Study of Model Components

To assess the contributions of individual HieraPTM components to PTM site prediction, we separately removed CNN-BiGRU, the center-aware Transformer, ESM-2-enhanced representations, and the complete three-dimensional structural branch and performed ablation experiments across eight PTM tasks. Macro-averaged performance is summarized in Table 4.

**Table 4.** Ablation study of the major components of HieraPTM across eight PTM prediction tasks.

| Model | AUROC | AUPRC | Accuracy<br>(%) | F1<br>(%) | Precision<br>(%) | Recall<br>(%) | Specificity<br>(%) | MCC |
|---|---|---|---|---|---|---|---|---|
| *Major-component ablations* |  |  |  |  |  |  |  |  |
| **HieraPTM (Full)** | **0.9237** | **0.8425** | 86.81 | **76.53** | 74.32 | 80.68 | 88.00 | **0.6685** |
| w/o CNN-BiGRU | 0.9087 | 0.8108 | 84.17 | 73.98 | 68.04 | 83.43 | 84.00 | 0.6290 |
| w/o Transformer | 0.9088 | 0.8089 | 85.47 | 75.04 | 70.30 | 81.33 | 85.63 | 0.6401 |
| w/o ESM-2 | 0.9055 | 0.8000 | 85.42 | 74.41 | 71.36 | 78.20 | 86.71 | 0.6301 |
| w/o 3D | 0.9217 | 0.8361 | **86.94** | 76.53 | **75.35** | 79.22 | **88.25** | 0.6673 |
| *Fusion-mechanism interventions* |  |  |  |  |  |  |  |  |
| Fixed ESM Gate | 0.9145 | 0.8236 | 85.69 | 74.33 | 73.54 | 78.12 | 87.47 | 0.6422 |
| w/o Center Global Access | 0.9200 | 0.8319 | 84.95 | 75.19 | 69.56 | **84.76** | 84.14 | 0.6495 |

Note: Values represent macro-averaged performance across eight PTM prediction tasks. Accuracy, F1, Precision, Recall, and Specificity are expressed as percentages. Bold values indicate the best performance for each metric based on unrounded results. The prefix “w/o” denotes removal of the corresponding component from HieraPTM.

Complete HieraPTM achieved the highest macro-averaged AUROC (0.9237) and AUPRC (0.8425), with F1 and MCC reaching 76.53% and 0.6685, respectively. Removing any module reduced both AUROC and AUPRC. ESM-2 made the most prominent contribution: its removal led to relative decreases of 1.97% and 5.04% in the two metrics, respectively, and reduced F1 from 76.53% to 74.41%. CNN-BiGRU and the Transformer also played important roles: their removal caused relative AUROC decreases of 1.62% and 1.62%, respectively, and relative AUPRC decreases of 3.76% and 3.99%, respectively. Notably, removing CNN-BiGRU increased Recall from 80.68% to 83.43%, but reduced Precision from 74.32% to 68.04% and MCC from 0.6685 to 0.6290. These results indicate that local sequence encoding, center-aware contextual integration, and pretrained protein representations all contribute substantially to overall discrimination.

In contrast, the overall gain from the three-dimensional structural branch was relatively limited. Removing this branch caused relative decreases in AUROC and AUPRC of 0.22% and 0.76%, respectively. Incorporating structural information increased Recall from 79.22% to 80.68%, but reduced Precision from 75.35% to 74.32%, while F1 remained approximately 76.53%. This indicates that three-dimensional structural information provides limited gains in ranking performance and introduces a further trade-off between classification sensitivity and precision.

### 3.5 Local Phosphosite Density Modulates Associations Between PTM Scores and Functional Annotations

To investigate potential links between protein PTM site prediction and phosphosite functional annotation status, we froze all parameters of the complete HieraPTM site prediction model and extracted phosphosite prediction scores. We combined these scores with the publicly available functional phosphosite annotations from FuncPhos-SEQ [16] for cross-task statistical analysis, focusing on three functional dimensions: molecular function, biological process, and protein interaction. To reduce the influence of overlap with upstream model training data, we included 57,795 ST/Y phosphosites that did not share identical 61-aa sequence windows with HieraPTM training samples. By integrating site prediction scores with functional annotations, we systematically assessed the statistical association between phosphorylation prediction information learned by HieraPTM and known functional annotations, providing a basis for further investigation of knowledge transfer from site prediction representations to functional phosphosite identification.

Using the phosphosite prediction scores produced by HieraPTM, we divided ST and Y sites separately into ten quantile groups and analyzed functional annotation distributions across score intervals. Functional annotations were markedly enriched among high-scoring sites, particularly in the highest decile, where molecular function annotation rates reached 14.2% for ST sites and 22.8% for Y sites (Supplementary Figure S4 A). This enrichment concentrated in the high-score range indicates that HieraPTM’s learned site prediction representations are associated not only with phosphorylation occurrence propensity but also with discriminative information concerning known functional annotation status.

To quantify this association while controlling for differences in functional annotation levels across proteins, we used within-protein conditional logistic regression, additionally adjusting for Ser/Thr residue type in the ST-site analysis. Phosphosite prediction scores were significantly positively associated with functional annotation status within proteins (Supplementary Figure S4 B). For each increase of one prespecified standard deviation in prediction score, the odds ratio for overall functional annotation was 1.250 (95% CI 1.177–1.328) for ST sites and 1.461 (95% CI 1.294–1.650) for Y sites. Further category-specific analyses found significant associations in the same direction for molecular function, biological process, and protein interaction, indicating consistency across multiple functional annotation dimensions.

Previous studies have shown that local multisite phosphorylation can regulate protein function through intersite dependencies and combinatorial effects [45], suggesting that phosphosite functional properties may depend on the local modification environment. We therefore further examined whether the local phosphorylation environment modulated these associations. Specifically, local phosphorylation density was quantified as the number of other observed phosphosites within 15 amino acids upstream and downstream of the target site. Interaction models were then used to assess differences in the association between prediction scores and functional annotations across density environments. The positive association was significantly stronger in high-density than in low-density environments (Supplementary Figure S4 C, D). Stratified functional annotation landscapes further showed enrichment of functional annotations in higher prediction-score ranges across local density conditions, while the strength of this association varied with the local phosphorylation environment.

These results demonstrate a significant within-protein statistical association between HieraPTM site prediction representations and functional phosphorylation annotations, modulated by the local phosphorylation environment. This finding is consistent with previously reported heterogeneity among phosphosites [15] and suggests that representations learned through multi-type PTM site prediction may support functional site prioritization.

Based on these findings, we further designed HieraFuncPhos (Figure 1) to transfer HieraPTM’s site recognition representations to functional phosphosite prediction. Through parameter-efficient functional adaptation and function-conditioned multiscale mechanical evidence retrieval, HieraFuncPhos integrates local and distal biophysical information, extending modification site recognition toward assessment of potential functional importance.

### 3.6 Predictive Performance against Established Functional Phosphosite Models

To evaluate how well HieraFuncPhos ranks phosphosites with curated regulatory or disease-association annotations above the sampled unlabeled background, we selected four representative functional phosphorylation prediction models as baselines: MMFuncPhos [19], FuncPhos-SEQ [16], FuncPhos-STR [18], and cDL-FuncPhos [17]. All models shared exactly the same training, validation, and independent test sets within each experimental round to ensure fair comparisons. Across nine runs, the unlabeled background samples were independently resampled, and the models were retrained on the corresponding run-specific datasets, while sharing the same pool of 9,007 positive phosphosites. We used AUROC and AUPRC as the primary evaluation metrics and additionally reported Accuracy, Precision, Recall, Specificity, F1-score, and MCC to comprehensively assess discrimination and classification performance. Final results were summarized as the mean and standard deviation across nine runs, reflecting performance variation under repeated background sampling and model retraining (Table 5). In addition, we report the complete results of all nine HieraFuncPhos replicates (Supplementary Table S6).

**Table 5.**
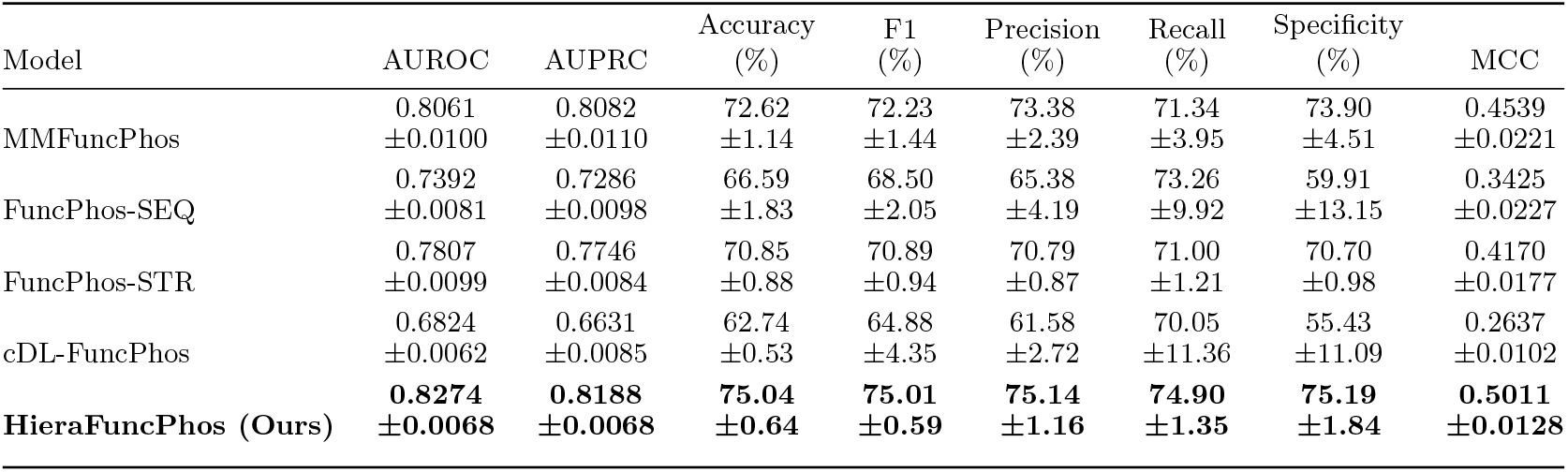
Performance comparison of HieraFuncPhos and baseline models. Results are reported as mean *±* standard deviation across nine runs with a shared positive pool, independently resampled unlabeled backgrounds, and separate model training. The standard deviation describes cross-run variation under this evaluation protocol. The best result for each metric is highlighted in bold.

| Model | AUROC | AUPRC | Accuracy<br>(%) | F1<br>(%) | Precision<br>(%) | Recall<br>(%) | Specificity<br>(%) | MCC |
|---|---|---|---|---|---|---|---|---|
| MMFuncPhos | 0.8061<br>±0.0100 | 0.8082<br>±0.0110 | 72.62<br>±1.14 | 72.23<br>±1.44 | 73.38<br>±2.39 | 71.34<br>±3.95 | 73.90<br>±4.51 | 0.4539<br>±0.0221 |
| FuncPhos-SEQ | 0.7392<br>±0.0081 | 0.7286<br>±0.0098 | 66.59<br>±1.83 | 68.50<br>±2.05 | 65.38<br>±4.19 | 73.26<br>±9.92 | 59.91<br>±13.15 | 0.3425<br>±0.0227 |
| FuncPhos-STR | 0.7807<br>±0.0099 | 0.7746<br>±0.0084 | 70.85<br>±0.88 | 70.89<br>±0.94 | 70.79<br>±0.87 | 71.00<br>±1.21 | 70.70<br>±0.98 | 0.4170<br>±0.0177 |
| cDL-FuncPhos | 0.6824<br>±0.0062 | 0.6631<br>±0.0085 | 62.74<br>±0.53 | 64.88<br>±4.35 | 61.58<br>±2.72 | 70.05<br>±11.36 | 55.43<br>±11.09 | 0.2637<br>±0.0102 |
| **HieraFuncPhos (Ours)** | **0.8274**<br>**±0.0068** | **0.8188**<br>**±0.0068** | **75.04**<br>**±0.64** | **75.01**<br>**±0.59** | **75.14**<br>**±1.16** | **74.90**<br>**±1.35** | **75.19**<br>**±1.84** | **0.5011**<br>**±0.0128** |

Overall, HieraFuncPhos achieved the highest mean performance on all eight evaluation metrics, demonstrating strong discrimination between phosphosites with qualifying annotations and sampled unlabeled background sites. HieraFuncPhos achieved an AUROC of 0.8274 *±* 0.0068 and an AUPRC of 0.8188 *±* 0.0068, outperforming the other models. The closest-performing method, MMFuncPhos, achieved AUROC and AUPRC values of 0.8061 and 0.8082, respectively. HieraFuncPhos improved these metrics by 0.0213 and 0.0106, corresponding to relative improvements of 2.64% and 1.31%. These results show that HieraFuncPhos more effectively ranks sites with qualifying annotations above sampled unlabeled sites on this benchmark.

HieraFuncPhos also demonstrated balanced advantages on threshold-dependent classification metrics. Its Accuracy, F1-score, and MCC reached 75.04%, 75.01%, and 0.5011, respectively, the highest values among all compared models. Compared with MMFuncPhos, Accuracy and F1-score improved by 2.42 and 2.78 percentage points, respectively, and MCC increased by an absolute 0.0472. Notably, HieraFuncPhos increased Recall from MMFuncPhos’s 71.34% to 74.90%, a gain of 3.56 percentage points. Meanwhile, Precision increased from 73.38% to 75.14%, and Specificity from 73.90% to 75.19%. These differences indicate improved recovery of sites with qualifying annotations and improved separation from the sampled background at the evaluation threshold.

To validate the effectiveness of key HieraFuncPhos components, we conducted stage-wise ablation experiments (Supplementary Tables S7, S8). Predictive performance improved continuously with the successive introduction of cross-task knowledge transfer, functional adaptation, and multiscale mechanical evidence correction, indicating complementary roles for these modules in functional phosphosite identification. Notably, analysis of the mechanical refinement module across different baseline prediction confidence levels revealed that its predictive gains increased substantially with baseline uncertainty, indicating that mechanical evidence provides greater complementary value for sites with ambiguous predictions.

To assess performance across distinct annotation categories, we further performed stratified analyses according to three functional annotation categories: molecular function-related, biological process-related, and protein interaction regulation (Supplementary Figure S5). HieraFuncPhos exhibited strong AUROC discrimination in all three categories and remained competitive on comprehensive metrics such as F1-score and MCC. This indicates that its predictive advantage is not restricted to a single functional annotation dimension but extends across the evaluated functional annotation categories.

Representative site analyses incorporating existing experimental studies showed that HieraFuncPhos maintained strong discrimination in low-structural-confidence regions and assigned high predicted probabilities to some known functional sites (Supplementary Figure S7). Among annotation-positive sites with pLDDT ¡ 50, 76 received predicted probabilities of at least 0.97. Specifically, ESR1 S118 (pLDDT = 30.81) and ESR2 Y36 (pLDDT = 38.78) received predicted probabilities of 0.973 and 0.978, respectively. GJA1 S262, S279, S365, and S373 also received high predicted probabilities of 0.969–0.974. The model also successfully identified MAP3K14 S817/S820 (0.794/0.784) and FANCD2 S331/S882 (0.710/0.638), which all four comparison methods failed to identify correctly; all these sites had pLDDT values below 35. Similar results were observed for SMN S63/S187 (0.697–0.877) and DEPTOR S244/S258/S297/S298 (0.638–0.896). These results indicate that HieraFuncPhos effectively identifies phosphosites with known functional annotations even when structural information has limited reliability, demonstrating prediction robustness in structurally uncertain regions.

### 3.7 Function-Guided Selection of Local and Distal Mechanical Evidence

To reveal how HieraFuncPhos converts hierarchical PTM recognition knowledge into multimodal representations of functional phosphosites, we analyzed representation organization and mechanical evidence integration across three stages: knowledge transfer, functional adaptation, and mechanical correction. UMAP visualization showed progressive reorganization of functional representations across stages (Figure 6A). The class separation metric (CSR) in the original high-dimensional space increased from 1.075 to 1.082 and reached 1.106 in the complete model. Further high-dimensional kNN analysis excluding neighbors from the same protein revealed distinct local neighborhood enrichment patterns for functional impact (FI), process regulation (PR), and protein interaction (PI) annotations. These patterns were reorganized through functional adaptation and mechanical correction (Figure 6B), indicating that hierarchical modeling enhances overall functional class separation and promotes local organization of function-related representations.

**Fig. 6.**
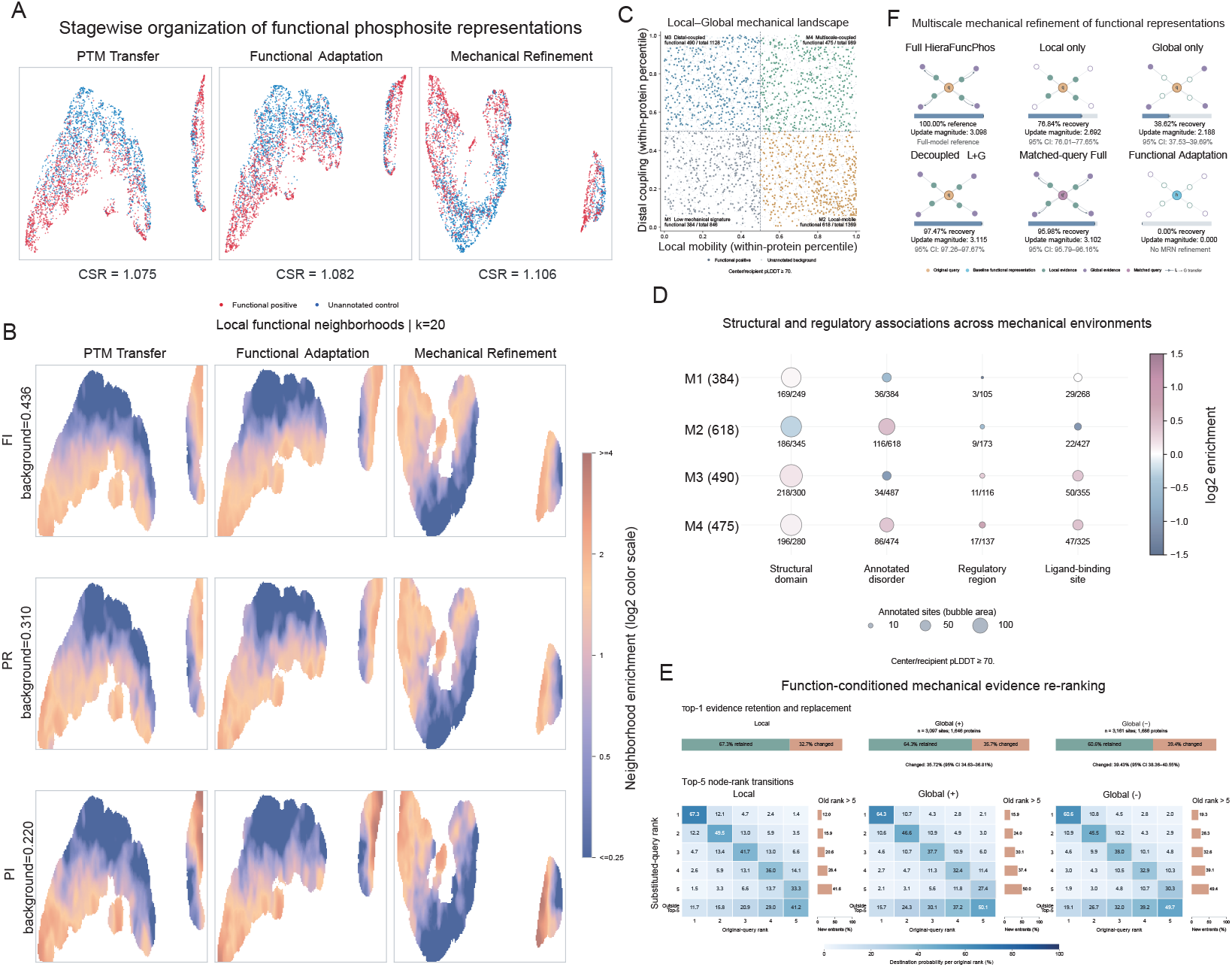
Adaptive Mechanical Evidence Retrieval and Functional Representation Refinement. **a**, UMAP visualization of functional representations across three stages, with CSR increasing from 1.075 to 1.106. **b**, Local neighborhood enrichment of FI, PR and PI annotations (*k* = 20). **c**, Four mechanical environments defined by local mobility and distal coupling. **d**, Enrichment of structural and regulatory annotations across mechanical environments. **e**, Functional-query substitution reorganizes local and distal residue rankings, with Top-1 replacement rates of 32.67%, 35.72% and 39.43% across three channels. **f**, Representation-update recovery under controlled mechanical readouts: Local-only (76.84%), Global-only (38.62%), decoupled Local + Global (97.47%) and matched-query Full (95.98%).

Mechanical environments also differed substantially among phosphosites. Based on within-protein standardized local mobility and distal mechanical coupling, evaluable sites were divided into low mechanical response (M1), local mobility (M2), distal coupling (M3), and multiscale coupling (M4) classes (Figure 6C). Structural and functional region enrichment analyses further showed that M2 was strongly associated with annotated disordered regions, whereas M3 and M4 exhibited different degrees of enrichment in domains, regulatory regions, and ligandbinding sites (Figure 6D). This association is consistent with previous evidence linking protein phosphorylation to intrinsically disordered regions [46]. These results reveal differences in the biophysical environments of functional phosphosites and provide biological context for site-adaptive integration of multiscale mechanical information.

A key feature of HieraFuncPhos is its use of functional representations to guide evidence retrieval from local and distal mechanical residues, rather than merely fusing precomputed dynamics descriptors. To quantitatively test this mechanism, we held candidate nodes, structures, and mechanical features fixed and separately replaced the functional queries in the Local, Global positive-coupling, and Global negative-coupling channels, examining changes in residue evidence priorities (Figure 6E). The mean Top-1 evidence replacement rates in the three channels reached 32.67%, 35.72%, and 39.43%, respectively, indicating that functional query conditioning can modulate the reading of both local and distal residues. Top-5 rank transition analysis showed that the originally fifth-ranked nodes dropped out of the top five after query replacement in 41.2%, 50.1%, and 49.7% of cases, respectively, while new candidate residues entered the high-priority evidence sets. These results show that functional representations participate in Local–Global scale integration and reorganize residue-level evidence priorities within fixed mechanical candidate sets. The Global branch additionally preserves information about distinct motion association patterns through positive- and negative-mechanical-coupling channels, providing direct computational support for function-conditioned dual-scale mechanical evidence retrieval.

Further controlled analyses showed that local and distal mechanical information jointly contribute to functional representation correction through complementary readings (Figure 6F). Relative to complete HieraFuncPhos, Local-only and Global-only recovered 76.84% and 38.62% of the full representation update, respectively. These results indicate that local mechanical evidence provides an important foundation for representation correction, with distal evidence offering complementary information. Independent reading and conditional integration of dual-scale evidence reproduce the great majority of the full representation update, indicating that function-conditioned retrieval and complementary multiscale reading jointly constitute important computational mechanisms of mechanical representation correction in HieraFuncPhos.

### 3.8 HieraFuncPhos Reveals Site-Specific Mechanical Responses of NTD Phosphorylation Sites in HSP90AA1

HSP90AA1 is an ATP-dependent molecular chaperone whose NTD H1 helix and lid region jointly participate in nucleotide-dependent conformational transitions. A recent solution NMR study [47] showed that, although T36 and T115 lie within the same functional regulatory unit, they regulate Hsp90 activity through distinct conformational dynamics pathways: T36E mainly alters conformational exchange in the ATP-analog-bound state, whereas T115E reshapes the NTD conformational energy landscape across multiple nucleotide states. Both phosphomimetic mutations are associated with inhibited ATPase activity, suggesting site-specific dynamic regulation within the same protein.

In test-set predictions, complete HieraFuncPhos assigned prediction scores of 0.7544 and 0.9457 to T36 and T115, respectively, both of which have independently reported regulatory roles. Using a shared wild-type reference structure, we performed a unified PRS analysis [48] of NTD residues 17–222 and found markedly different mechanical response profiles for the two sites. The mechanical response of T36 was concentrated mainly in the local region, with distal PRS enrichment of only 0.07. In contrast, T115 showed a stronger response in the sequence-distal H1 region, with distal PRS enrichment reaching 1.47.

Consistent with this mechanical environmental heterogeneity, conditional mechanical integration in Hiera-FuncPhos was strongly site-dependent. For T36, the model allocated 81.67% of its internal routing weight to the Local branch and only 18.08% to the Global branch, indicating a pronounced preference for local evidence. In contrast, the Local and Global weights for T115 were 53.82% and 45.68%, respectively. Its Global weight was 27.60 percentage points higher than that of T36, producing a more balanced pattern of multiscale evidence integration. This adaptive routing trend was consistent with the independently computed spatial distribution of mechanical responses, indicating that HieraFuncPhos develops distinct strategies for reading local and distal mechanical evidence according to site environments within the same protein rather than using a uniform feature fusion ratio.

To further assess the experimental relevance of these computational mechanical responses, we compared residue-level PRS with published NMR data for the T36E and T115E phosphomimetic mutants (Figure 7). Backbone and methyl chemical shift perturbation analyses showed that the principal response to T36E was concentrated near T36, whereas T115E induced broader changes in chemical environments across H1, the lid, and associated regions. These experimental observations show qualitative regional agreement with the computed distinction between predominantly local mechanical responses at T36 and stronger distal responses at T115.

**Fig. 7.**
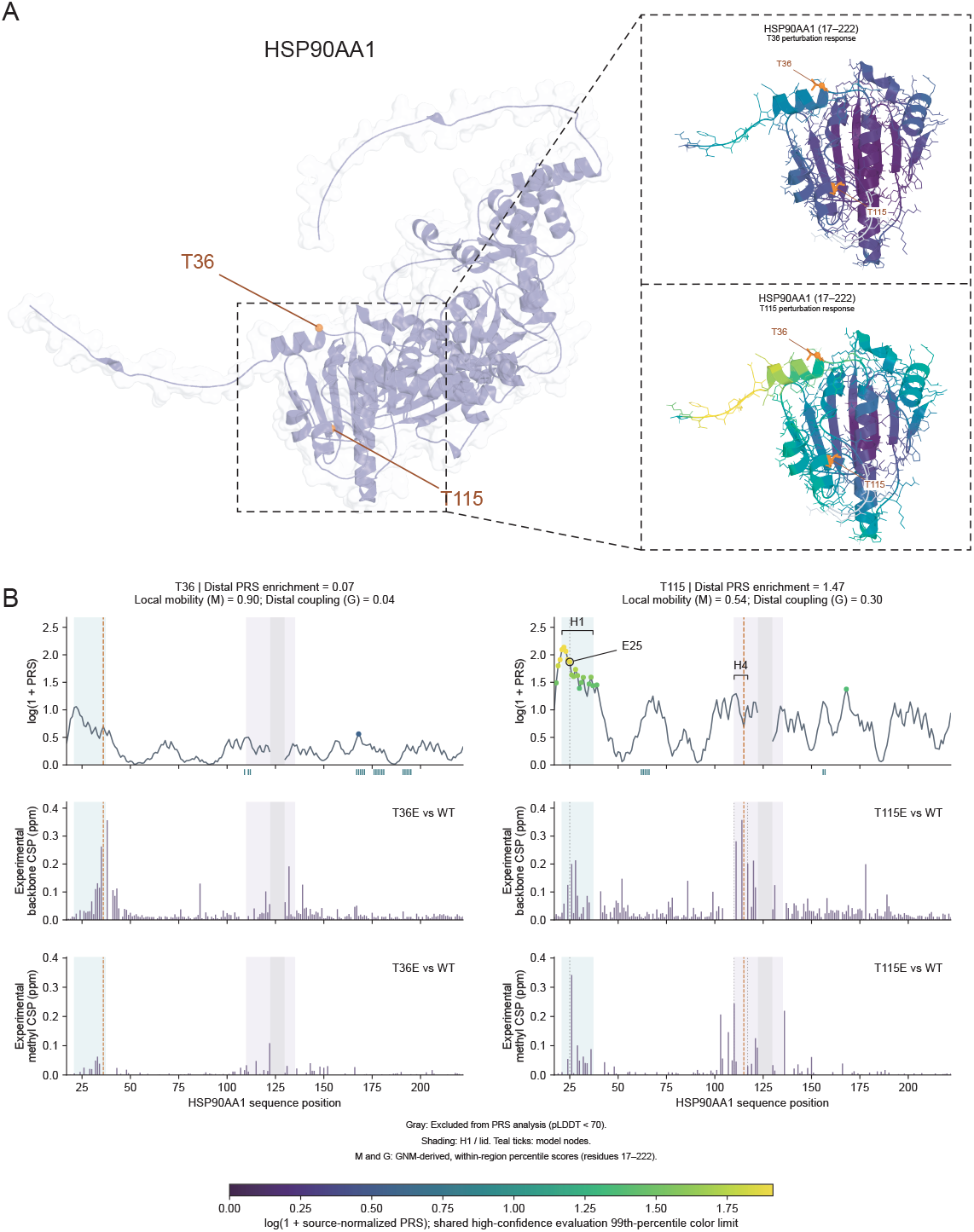
Site-specific reference-structure mechanical response patterns of HSP90AA1 phosphorylation sites compared with experimental NMR observations. **a**, HSP90AA1 structure highlighting T36 and T115, with their predicted perturbation-response scanning (PRS) patterns in the N-terminal domain. **b**, Residue-level PRS profiles for T36 (left) and T115 (right), compared with experimental backbone and methyl chemical shift perturbations (CSPs) of T36E and T115E. T36 exhibits predominantly local responses, whereas T115 shows stronger distal responses involving the H1 region. Shading indicates H1 and lid regions; gray denotes residues excluded from PRS analysis (pLDDT *<* 70).

Notably, an independent NMR structural study found that the negative charge introduced by T115E can cause electrostatic repulsion with E25 in the H1 region, accompanied by an approximately 30° conformational rotation of the lid-region H4 helix and associated structural rearrangements. Reference-structure PRS analysis in HieraFuncPhos also identified H1 as a major high-response region for T115, showing qualitative correspondence between computed mechanical sensitivity and an experimentally implicated regulatory region. In contrast, T36E remained relatively stable in its overall structure, with its principal regulatory effect manifested as a selective change in conformational exchange during ATP binding. This is consistent with the more restricted reference-structure response fingerprint of T36.

The comparison of test-set predictions with experimental cases indicates that complete HieraFuncPhos identifies two phosphosites with known regulatory roles and reveals heterogeneous mechanical responses associated with different sites within the same protein. Within the same protein, the local mechanical response of T36 corresponds to the model’s preference for Local reading, whereas the pronounced distal H1 response of T115 is accompanied by a higher proportion of Global integration. Together with independent NMR measurements and structural mechanistic evidence, these results provide a biophysically meaningful case supporting function-conditioned multi-scale mechanical integration in HieraFuncPhos and demonstrate its ability to provide site-specific regulatory clues beyond functional site identification.

## 4 Discussion and conclusion

This study introduces the HieraPTM–HieraFuncPhos hierarchical deep learning framework to address effective integration of multiscale biological information in PTM site prediction and establish a knowledge link between multi-type PTM site recognition and functional phosphosite prediction. Rather than modeling the two tasks independently, the framework combines hierarchical representation learning and cross-task knowledge transfer to improve predictive performance while examining the distinct roles of biological evidence at different scales in modification recognition and functional importance assessment. Systematic benchmarking showed competitive performance on both prediction tasks, with most evaluation metrics exceeding those of existing comparison methods, demonstrating good predictive ability and generalization potential. Ablation experiments further validated the hierarchical hybrid architecture, showing that local sequence modeling, protein-context fusion, and three-dimensional structural correction play complementary roles at different information levels. Their progressive integration and joint optimization improve overall discrimination.

For PTM site recognition, HieraPTM hierarchically integrates local sequence, protein context, and three-dimensional microenvironments through center-aware contextual modeling, fine-grained residual gating, and structure-guided incremental correction. Systematic ablation and interpretability analyses showed that center-aware modeling helps strengthen PTM-specific residue–position recognition patterns, protein language models mainly supplement sites with ambiguous local evidence, and three-dimensional structural information provides more localized discriminative corrections. These findings indicate that the modalities provide distinct discriminative contributions and complementary information rather than simply interchangeable features, offering new insight into how multiscale biological information jointly supports PTM site recognition.

For functional phosphosite prediction, HieraFuncPhos transfers PTM site recognition knowledge to annotation-based discrimination through LoRA-based functional adaptation and selectively integrates local and distal mechanical information through function-conditioned mechanical evidence retrieval. The functional phosphosite prediction results further indicate the potential value of transferring representations learned from multi-type PTM site recognition to the discrimination of phosphosites with curated functional annotations, while site-specific biophysical environments provide task-specific information that complements shared recognition knowledge.

The value of this study lies in connecting high-quality PTM data, multiscale biological information integration, and functional annotation-based phosphosite prediction. The framework improves PTM site recognition and the discrimination of phosphosites with curated functional annotations from those lacking such annotations, providing an approach to hierarchical information integration and cross-task knowledge transfer for multimodal protein analysis. The modification recognition patterns and structural–mechanical features associated with functional annotations identified by the models may also provide computational guidance for prioritizing disease-associated regulatory site candidates, identifying potential intervention targets for further investigation, and exploring protein regulatory mechanisms. However, the absence of functional annotations does not necessarily indicate a lack of biological function, and incomplete annotation coverage may influence model predictions. Moreover, predicted structures and elastic network models cannot fully represent the dynamic changes induced by actual phosphorylation events. Generalization of annotation-based predictions across protein families also requires further validation. Future studies could integrate structural dynamics analyses of modified protein states with experimental functional measurements to further investigate the biological relevance of cross-task knowledge transfer and structural–mechanical evidence. Extending the framework to other PTM types with available functional annotations may provide a foundation for more general, accurate, and biologically interpretable protein modification analysis.

## Supporting information

This is the supplementary material.

## 6 Data and code availability

The source code and associated dataset splits for HieraPTM and HieraFuncPhos are publicly available at GitHub (https://github.com/yuan86927/HieraPTM-HieraFuncPhos). The repository provides model implementations, training and evaluation scripts, hyperparameter configurations, and dataset splits for eight PTM-site prediction tasks and nine functional phosphorylation prediction replicates.

## 7 Author contributions statement

Yuzhi Yuan: Writing – original draft, Visualization, Validation, Methodology, Software, Investigation, Formal analysis, Data curation, Conceptualization; Chang Ye (Corresponding author): Writing – review & editing, Supervision, Project administration, Conceptualization, Methodology, Resources; Tianchi Lu (Corresponding author): Writing – review & editing, Supervision, Project administration, Conceptualization, Methodology, Resources; Di Wang: Supervision, Writing – review & editing; Pengyu Chen: Supervision, Writing – review & editing; Yanze Niu: Supervision, Writing – review & editing; Zhiyuan Yang: Writing – review & editing. Hengjia Zhang: Writing – review & editing. Xicheng He: Investigation, Validation, Writing – review & editing. Hao Li: Data curation, Formal analysis, Validation. Ziwei Su: Visualization, Data curation, Writing – review & editing. Changwei Lin: Methodology, Software, Formal analysis. Tianhang Chen: Methodology, Validation, Writing – review & editing.

## 8 Acknowledgments

We thank all members of our research group for their valuable discussions and feedback throughout this project.

## References

[1] Yamanashi, Y., Kanai, M.: Catalytic strategies for post-translational modifications regulating protein higher-order structure and properties. Chemical Society Reviews 55(8), 4409–4425 (2026) 10.1039/d6cs00107f

[2] Müller, J., Kuster, B., The, M.: Computational approaches for pathway-centric analysis of protein post-translational modifications. PROTEOMICS 25(21-22), 41–61 (2025) 10.1002/pmic.70055 https://analyticalsciencejournals.onlinelibrary.wiley.com/doi/pdf/10.1002/pmic.70055

[3] Beltrao, P., Albanèse, V., Kenner, L.R., Swaney, D.L., Burlingame, A., Villén, J., Lim, W.A., Fraser, J.S., Frydman, J., Krogan, N.J.: Systematic functional prioritization of protein posttranslational modifications.Cell 150(2), 413–425 (2012) 10.1016/j.cell.2012.05.036

[4] Zhang, S., Li, J., Chen, M., Huang, H.: Post-translational modifications in proteins: Prediction methods, biological functions, and diseases. MedComm 7(4), 70729 (2026) 10.1002/mco2.70729 https://onlinelibrary.wiley.com/doi/pdf/10.1002/mco2.70729

[5] Ubersax, J.A., Ferrell Jr, J.E.: Mechanisms of specificity in protein phosphorylation. Nature Reviews Molecular Cell Biology 8(7), 530–541 (2007) 10.1038/nrm2203

[6] Houles, T., Yoon, S.-O., Roux, P.P.: The expanding landscape of canonical and non-canonical protein phosphorylation. Trends in Biochemical Sciences 49(11), 986–999 (2024) https://doi.org/10.1016/j.tibs.2024.08.004. doi: 10.1016/j.tibs.2024.08.004

[7] Wen, B., Wang, C., Li, K., Han, P., Holt, M.V., Savage, S.R., Lei, J.T., Dou, Y., Shi, Z., Li, Y., Zhang, B.: Deepmvp: deep learning models trained on high-quality data accurately predict PTM sites and variant-induced alterations. Nature Methods 22(9), 1857–1867 (2025) 10.1038/s41592-025-02797-x

[8] Rives, A., Meier, J., Sercu, T., Goyal, S., Lin, Z., Liu, J., Guo, D., Ott, M., Zitnick, C.L., Ma, J., Fergus, R.: Biological structure and function emerge from scaling unsupervised learning to 250 million protein sequences. Proceedings of the National Academy of Sciences 118(15), 2016239118 (2021) https://doi.org/10.1073/pnas.2016239118. doi: 10.1073/pnas.2016239118

[9] Brandes, N., Ofer, D., Peleg, Y., Rappoport, N., Linial, M.: ProteinBERT: a universal deep-learning model of protein sequence and function. Bioinformatics 38(8), 2102–2110 (2022) 10.1093/bioinformatics/btac020

[10] Xu, Z., Zhong, H., He, B., Wang, X., Lu, T.: PTransIPs: Identification of phosphorylation sites enhanced by protein PLM embeddings. IEEE Journal of Biomedical and Health Informatics 28(6), 3762–3771 (2024) 10.1109/JBHI.2024.3377362

[11] Zhai, J., Wang, Z., Tang, C., Zhong, H., Xu, Z., Liu, Y., Xu, S., Wang, J., Huang, D., Lu, T.: A general language model for peptide function identification. https://arxiv.org/abs/2502.15610 (2025)

[12] Li, Z., Li, M., Zhu, L., Zhang, W.: Improving PTM site prediction by coupling of multi-granularity structure and multi-scale sequence representation. Proceedings of the AAAI Conference on Artificial Intelligence 38(1), 188–196 (2024) 10.1609/aaai.v38i1.27770

[13] Wang, C., Jiang, Q., Yu, D., Guan, J., Chen, Z., Lu, X., Yan, B., Qin, J., Liu, Y., Wang, J.: Deepptmpred: a multi-modal deep learning framework for accurate prediction of protein post-translational modification sites. Briefings in Bioinformatics 27(3), 321 (2026) 10.1093/bib/bbag321 https://academic.oup.com/bib/article-pdf/27/3/bbag321/68542133/bbag321.pdf

[14] Peng, F.Z., Wang, C., Chen, T., Schussheim, B., Vincoff, S., Chatterjee, P.: PTM-Mamba: a PTM-aware protein language model with bidirectional gated mamba blocks. Nature Methods 22(5), 945–949 (2025) 10.1038/s41592-025-02656-9

[15] Ochoa, D., Jarnuczak, A.F., Viéitez, C., Gehre, M., Soucheray, M., Mateus, A., Kleefeldt, A.A., Hill, A., Garcia-Alonso, L., Stein, F., Krogan, N.J., Savitski, M.M., Swaney, D.L., Vizcaíno, J.A., Noh, K.-M., Beltrao, P.: The functional landscape of the human phosphoproteome. Nature Biotechnology 38(3), 365–373 (2020) 10.1038/s41587-019-0344-3

[16] Liang, Z., Liu, T., Li, Q., Zhang, G., Zhang, B., Du, X., Liu, J., Chen, Z., Ding, H., Hu, G., Lin, H., Zhu, F., Luo, C.: Deciphering the functional landscape of phosphosites with deep neural network. Cell Reports 42(9) (2023) https://doi.org/10.1016/j.celrep.2023.113048. doi: 10.1016/j.celrep.2023.113048

[17] Zhu, F., Yang, S., Meng, F., Zheng, Y., Ku, X., Luo, C., Hu, G., Liang, Z.: Leveraging protein dynamics to identify functional phosphorylation sites using deep learning models. Journal of Chemical Information and Modeling 62(14), 3331–3345 (2022) 10.1021/acs.jcim.2c00484 https://pubs.acs.org/jcisd8/ar ticle-pdf/62/14/3331/8905020/ci2c00484.pdf

[18] Zhang, G., Zhang, C., Cai, M., Luo, C., Zhu, F., Liang, Z.: Funcphos-str: An integrated deep neural network for functional phosphosite prediction based on alphafold protein structure and dynamics. International Journal of Biological Macromolecules 266, 131180 (2024) 10.1016/j.ijbiomac.2024.131180

[19] Xie, J., Dong, R., Zhu, J., Lin, H., Wang, S., Lai, L.: Mmfuncphos: A multi-modal learning framework for identifying functional phosphorylation sites and their regulatory types. Advanced Science 12(9), 2410981 (2025) 10.1002/advs.202410981 https://advanced.onlinelibrary.wiley.com/doi/pdf/10.1002/advs.202410981

[20] Hong, X., Lv, J., Li, Z., Zhu, J., Li, J., Rahman, M.U., Wei, T., Mu, J., Chen, H.-F.: Deep learning model of post-translational modification regulating liquid-liquid phase separation. Communications Chemistry 8(1), 393 (2025) 10.1038/s42004-025-01773-y

[21] Suskiewicz, M.J.: The logic of protein post-translational modifications (ptms): Chemistry, mechanisms and evolution of protein regulation through covalent attachments. BioEssays 46(3), 2300178 (2024) 10.1002/bies.202300178 https://onlinelibrary.wiley.com/doi/pdf/10.1002/bies.202300178

[22] Hu, E.J., Shen, Y., Wallis, P., Allen-Zhu, Z., Li, Y., Wang, S., Wang, L., Chen, W.: LoRA: Low-Rank Adaptation of Large Language Models. In: International Conference on Learning Representations (2022). https://mlanthology.org/iclr/2022/hu2022iclr-lora/

[23] The UniProt Consortium: Uniprot: the universal protein knowledgebase in 2023. Nucleic Acids Research 51(D1), 523–531 (2023) 10.1093/nar/gkac1052

[24] Hornbeck, P.V., Zhang, B., Murray, B., Kornhauser, J.M., Latham, V., Skrzypek, E.: PhosphoSitePlus, 2014: mutations, PTMs and recalibrations. Nucleic Acids Research 43(D1), 512–520 (2015) 10.1093/nar/gku1267

[25] Lin, Z., Akin, H., Rao, R., Hie, B., Zhu, Z., Lu, W., Smetanin, N., Verkuil, R., Kabeli, O., Shmueli, Y., Santos Costa, A., Fazel-Zarandi, M., Sercu, T., Candido, S., Rives, A.: Evolutionary-scale prediction of atomic-level protein structure with a language model. Science 379(6637), 1123–1130 (2023) https://doi.org/10.1126/science.ade2574. doi: 10.1126/science.ade2574

[26] Jumper, J., Evans, R., Pritzel, A., Green, T., Figurnov, M., Ronneberger, O., Tunyasuvunakool, K., Bates, R., Žídek, A., Potapenko, A., Bridgland, A., Meyer, C., Kohl, S.A.A., Ballard, A.J., Cowie, A., Romera-Paredes, B., Nikolov, S., Jain, R., Adler, J., Back, T., Petersen, S., Reiman, D., Clancy, E., Zielinski, M., Steinegger, M., Pacholska, M., Berghammer, T., Bodenstein, S., Silver, D., Vinyals, O., Senior, A.W., Kavukcuoglu, K., Kohli, P., Hassabis, D.: Highly accurate protein structure prediction with alphafold. Nature 596(7873), 583–589 (2021) 10.1038/s41586-021-03819-2

[27] Varadi, M., Bertoni, D., Magana, P., Paramval, U., Pidruchna, I., Radhakrishnan, M., Tsenkov, M., Nair, S., Mirdita, M., Yeo, J., Kovalevskiy, O., Tunyasuvunakool, K., Laydon, A., Žídek, A., Tomlinson, H., Hariharan, D., Abrahamson, J., Green, T., Jumper, J., Birney, E., Steinegger, M., Hassabis, D., Velankar, S.: Alphafold protein structure database in 2024: providing structure coverage for over 214 million protein sequences. Nucleic Acids Research 52(D1), 368–375 (2024) 10.1093/nar/gkad1011

[28] Fu, L., Niu, B., Zhu, Z., Wu, S., Li, W.: Cd-hit: accelerated for clustering the next-generation sequencing data. Bioinformatics 28(23), 3150–3152 (2012) 10.1093/bioinformatics/bts565 https://academic.oup.com/bioinformatics/article-pdf/28/23/3150/50695216/bioinformatics28233150.pdf

[29] Cho, K., Merriënboer, B., Gulcehre, C., Bahdanau, D., Bougares, F., Schwenk, H., Bengio, Y.: Learning phrase representations using RNN encoder–decoder for statistical machine translation. In: Moschitti, A., Pang, B., Daelemans, W. (eds.) Proceedings of the 2014 Conference on Empirical Methods in Natural Language Processing (EMNLP), pp. 1724–1734. Association for Computational Linguistics, Doha, Qatar (2014). 10.3115/v1/D14-1179. https://aclanthology.org/D14-1179/

[30] Vaswani, A., Shazeer, N., Parmar, N., Uszkoreit, J., Jones, L., Gomez, A.N., Kaiser, L., Polosukhin, I.: Attention is all you need. In: Proceedings of the 31st International Conference on Neural Information Processing Systems. NIPS’17, pp. 6000–6010. Curran Associates Inc., Red Hook, NY, USA (2017)

[31] Veličković, P., Cucurull, G., Casanova, A., Romero, A., Liò, P., Bengio, Y.: Graph Attention Networks. In: International Conference on Learning Representations (2018). https://mlanthology.org/iclr/2018/velickovic2018iclr-graph/

[32] Gutierrez, C.S., Kassim, A.A., Gutierrez, B.D., Raines, R.T.: Sitetack: a deep learning model that improves ptm prediction by using known ptms. Bioinformatics 40(11), 602 (2024) 10.1093/bioinformatics/btae602 https://academic.oup.com/bioinformatics/article-pdf/40/11/btae602/60592587/btae602.pdf

[33] Yang, H., Wang, M., Liu, X., Zhao, X.-M., Li, A.: PhosIDN: an integrated deep neural network for improving protein phosphorylation site prediction by combining sequence and protein-protein interaction information. Bioinformatics (Oxford, England) 37(24), 4668–4676 (2021) 10.1093/bioinformatics/btab551

[34] Zhang, J., Cao, H., Gao, Z., Wang, Y., Li, S., Xu, J., Tan, C., Zhu, J., Hsieh, C.-Y., Gu, C., Heng, P.A.: Learning the PTM code through a coarse-to-fine mechanism-aware framework. Nature Communications 17(1), 6450 (2026) 10.1038/s41467-026-73148-3

[35] Zhou, H., Tan, W., Shi, S.: Deepgpgs: a novel deep learning framework for predicting arginine methylation sites combined with gaussian prior and gated self-attention mechanism. Briefings in Bioinformatics 24(2), 018 (2023) 10.1093/bib/bbad018 https://academic.oup.com/bib/article-pdf/24/2/bbad018/4956 0476/bbad018.pdf

[36] Li, Z., Li, S., Luo, M., Jhong, J.-H., Li, W., Yao, L., Pang, Y., Wang, Z., Wang, R., Ma, R., Yu, J., Huang, Y., Zhu, X., Cheng, Q., Feng, H., Zhang, J., Wang, C., Hsu, J.-K., Chang, W.-C., Wei, F.-X., Huang, H.-D., Lee, T.-Y.: dbptm in 2022: an updated database for exploring regulatory networks and functional associations of protein post-translational modifications. Nucleic Acids Research 50(D1), 471–479 (2022) 10.1093/nar/gkab1017 https://academic.oup.com/nar/article-pdf/50/D1/D471/42057610/gkab1017.pdf

[37] Liu, Y., Wang, X., Zhong, H., Zhai, J., Gong, X., Lu, T.: Phosf3c: a feature fusion architecture with finetuned protein language model and conformer for prediction of general phosphorylation site. Briefings in Bioinformatics 26(3), 242 (2025) 10.1093/bib/bbaf242 https://academic.oup.com/bib/article-pdf/26/3/bbaf242/63368372/bbaf242.pdf

[38] McInnes, L., Healy, J., Saul, N., Großberger, L.: Umap: Uniform manifold approximation and projection. Journal of Open Source Software 3(29), 861 (2018) 10.21105/joss.00861

[39] Schwarz, F., Aebi, M.: Mechanisms and principles of n-linked protein glycosylation. Current Opinion in Structural Biology 21(5), 576–582 (2011) 10.1016/j.sbi.2011.08.005.Carbohydrates and glycoconjugates/Biophysical methods

[40] Gareau, J.R., Lima, C.D.: The SUMO pathway: emerging mechanisms that shape specificity, conjugation and recognition. Nature Reviews Molecular Cell Biology 11(12), 861–871 (2010) 10.1038/nrm3011

[41] Lu, K.P., Zhou, X.Z.: The prolyl isomerase pin1: a pivotal new twist in phosphorylation signalling and disease. Nature Reviews Molecular Cell Biology 8(11), 904–916 (2007) 10.1038/nrm2261

[42] Han, J., Fang, Z., Han, B., Ye, B., Lin, W., Jiang, Y., Han, X., Wang, X., Wu, G., Wang, Y., Liang, G.: Deubiquitinase josd2 improves calcium handling and attenuates cardiac hypertrophy and dysfunction by stabilizing serca2a in cardiomyocytes. Nature Cardiovascular Research 2(8), 764–777 (2023) 10.1038/s44161-023-00313-y

[43] Chen, W., Smeekens, J.M., Wu, R.: Systematic and site-specific analysis of n-sialoglycosylated proteins on the cell surface by integrating click chemistry and MS-based proteomics. Chemical Science 6(8), 4681–4689 (2015) 10.1039/c5sc01124h

[44] ProteomeXchange Consortium: PXD028593: (Phospho)-proteomic characterization of longitudinal biopsies from HCC patients treated with sorafenib. PRIDE repository. ProteomeXchange dataset PXD028593 (2026)

[45] Conti, M.M., Li, R., Narvàez Ramos, M.A., Zhu, L.J., Fazzio, T.G., Benanti, J.A.: Phosphosite scanning reveals a complex phosphorylation code underlying CDK-dependent activation of hcm1. Nature Communications 14(1), 310 (2023) 10.1038/s41467-023-36035-9

[46] Iakoucheva, L.M., Radivojac, P., Brown, C.J., O’Connor, T.R., Sikes, J.G., Obradovic, Z., Dunker, A.K.: The importance of intrinsic disorder for protein phosphorylation. Nucleic Acids Research 32(3), 1037–1049 (2004) 10.1093/nar/gkh253

[47] Wan, C., Zhu, L., Wang, S., Zhang, X., Zhao, J., Qu, X., Huang, C.: Distinct phosphorylation mechanisms as dynamic switches for hsp90 regulation. Nature Communications 17(1), 6577 (2026) 10.1038/s41467-026-73400-w

[48] Atilgan, C., Gerek, Z.N., Ozkan, S.B., Atilgan, A.R.: Manipulation of conformational change in proteins by single-residue perturbations. Biophysical Journal 99(3), 933–943 (2010) 10.1016/j.bpj.2010.05.020

