## Supplementary material for "HieraPTM: From PTM Site Recognition to Functional Phosphosite Prediction through Hierarchical Context and Protein Mechanics": This is the supplementary material.

### 1 Supplementary Method

#### Datasets and preprocessing

This study proposes HieraFuncPhos, a framework for functional phosphorylation site prediction that combines hierarchical PTM knowledge transfer with multiscale structural-dynamics adaptation. We constructed a benchmark for functional phosphorylation site prediction using experimentally identified serine, threonine, and tyrosine (STY) phosphosites from the FuncPhos-STR (FuncSTR) dataset, directly adopting the functional annotation labels provided in the original dataset. Sites with existing functional annotations were designated as positive samples, whereas sites lacking such annotations were treated as unlabeled background. To evaluate predictive performance under different background sampling conditions, we conducted nine runs (Rep1–Rep9) with independently resampled unlabeled backgrounds and separate model training. Specifically, the same pool of 9,007 annotated positive sites was used across all runs, while 9,007 background sites were independently sampled from the unlabeled candidate pool in each run, yielding nine balanced datasets with 18,014 sites each. Each dataset was partitioned by stratified sampling into a model development set (80%, 14,410 sites) and an independent test set (20%, 3,604 sites). First, five-fold cross-validation was performed on the development set for model development, hyperparameter optimization, and model configuration selection. After the optimal configuration was determined, the final model was retrained using a fixed training–validation split within the development set, with early stopping and weight selection based on validation performance, followed by a uniform evaluation on the independent test set. Throughout this process, the independent test set was not involved in model training, hyperparameter selection, or early-stopping decisions.

For each site, sequence, structure, and dynamics inputs were aligned according to a unified protein sequence and residue numbering. All features were linked through a unique site identifier, and missing or unavailable inputs were recorded using explicit masks; sites were not directly removed solely because of low structural confidence.

#### Hierarchical PTM knowledge transfer

HieraFuncPhos uses HieraPTM pretrained on Task1 as its representation backbone, transferring the local sequence grammar, protein context, and static three-dimensional structural knowledge learned from PTM-site occurrence recognition to the functional prediction task. According to the acceptor residue type, ST sites and Y sites use the corresponding phosphorylation-ST and phosphorylation-Y pretrained encoders, respectively. The two encoders have separate parameters, but STY sites are trained jointly during functional adaptation, with a shared downstream functional fusion module and binary classification head.

For a target site $i$, a 61-residue sequence window centered on that site is used, integrating frozen ESM-2 representations and local static structural inputs. The local sequence and static graph nodes in the backbone use frozen ESM-2 150M representations, whereas the protein-wide context uses frozen ESM-2 3B representations.

To achieve parameter-efficient functional adaptation, low-rank adaptation (LoRA) is introduced into the center-aware Transformer. For a pretrained linear mapping $W$, its effective weight update is expressed as

$$W' = W + \frac{\alpha}{r}BA, \tag{S1}$$

where $A \in \mathbb{R}^{r\times d_{\text{in}}}$, $B \in \mathbb{R}^{d_{\text{out}}\times r}$, and $r$ is the low-rank dimension. The original weights $W$ remain frozen, and $B$ is initialized to zero, so that the low-rank module does not alter the pretrained linear mapping at initialization. The attention Q, V, and output mappings use $r = 8$, and the two linear mappings in the FFN use $r = 4$, with $\alpha = 2r$ in all cases. In addition to the low-rank parameters, the pretrained ESM fusion gate and structural interaction scaling parameters also participate in functional adaptation; the original sequence encoding network, the main Transformer weights, and the graph encoding network weights remain frozen.

Let the site representation output by the residue-matched encoder be $s_i \in \mathbb{R}^{192}$, and the protein-wide representation provided by frozen ESM-2 3B be $e_i^G \in \mathbb{R}^{2560}$. Projection, gated fusion, and functional residual adaptation yield the functional representation

$$\begin{aligned} u_i &= P_G(e_i^G), \\ g_i^G &= \sigma\left(W_G[s_i \| u_i] + b_G\right), \\ v_i &= s_i + g_i^G u_i, \\ h_i^F &= v_i + R_F(v_i), \end{aligned} \tag{S2}$$

where $P_G$ projects the protein-wide representation to 192 dimensions, $g_i^G$ is a site-dependent scalar gate, $R_F$ is the functional residual adaptation module, and $\|$ denotes vector concatenation. The base functional predictor produces the following outputs through a newly constructed shared classification head:

$$z_i^F = H_F(h_i^F), \qquad p_i^F = \sigma(z_i^F), \tag{S3}$$

where $H_F$ is the functional classification head, and $\sigma$ denotes the sigmoid function.

#### Multiscale structural-dynamics representation

To supplement mechanical properties that are difficult to describe explicitly using static structures, an elastic network is constructed from the structural coordinates in AlphaFold Database v6, combining the Gaussian network model (GNM) and the anisotropic network model (ANM) to extract three types of dynamics evidence: Motion, Response, and Network. Motion describes residue flexibility, multiscale motion correlations, and their stability; Response describes perturbation-response scanning (PRS) under the low-frequency ANM approximation and the relative dynamic flexibility index (DFI); Network describes motion communities, the organization of node connectivity, and cross-region connections.

The dynamics inputs are further divided into two scales: Local and Global. The local neighborhood contains the center site and up to 24 neighboring residues, with a total of 25 node slots retained. Local candidates have a three-dimensional distance from the center of no more than 12 Å, or a sequence distance of no more than 15 residues. Distal candidates must simultaneously have a sequence distance greater than 30 residues and a three-dimensional distance greater than 12 Å, with nodes already included in the Task1 static graph excluded. Distal nodes are ranked according to multiscale GNM correlations, sign consistency, and the correlation increment relative to the geometric background, retaining up to 24 positively coupled nodes and 24 negatively coupled nodes, respectively. All candidate selection and dynamics feature calculations are completed before training, without using functional labels.

For site $i$ and its candidate node $j$, the three types of physical features are denoted by $x_{ij}^{(M)}$, $x_{ij}^{(R)}$, and $x_{ij}^{(N)}$, with dimensions of 10, 8, and 10, respectively. Each view is mapped to a shared 128-dimensional space through an independent projection:

$$v_{ij}^{(k)} = \phi_k\left(x_{ij}^{(k)}\right), \qquad k \in \{M, R, N\}. \tag{S4}$$

Conditioned view weights are then computed based on the functional representation, node quality, and view and scale information:

$$\beta_{ij}^{(k)} = \mathrm{MaskedSoftmax}_k\left[a_\theta\left(v_{ij}^{(k)}, h_i^F, q_{ij}, e_k, e_{\mathrm{scale}}\right)\right], \tag{S5}$$

where $q_{ij}$ denotes node structural quality information, and $e_k$ and $e_{\mathrm{scale}}$ are the view and scale embeddings, respectively. MaskedSoftmax excludes unavailable views and renormalizes the remaining views.

The physical evidence is further fused with frozen ESM-2 3B node semantic representations to obtain the dynamics node representation

$$t_{ij} = m_{ij}\,\mathrm{LN}\left[P_E\left(e_{ij}^{\mathrm{ESM}}\right) + \sum_{k \in \{M,R,N\}} \beta_{ij}^{(k)} v_{ij}^{(k)}\right], \tag{S6}$$

where $m_{ij}$ is the node validity mask, and $P_E$ maps the node semantic representations to 128 dimensions. Normalization parameters for the physical features are estimated only from the training data in the current fold and applied to the corresponding validation and test data.

#### Function-conditioned mechanical refinement

The local branch uses two layers of graph attention with physical edge biases to encode the mechanical microenvironment of the center site; the global branch separately retrieves positively and negatively coupled distal motion evidence. Both scales are conditioned on the base functional representation $h_i^F$, rather than constructing functional classifiers independently.

To capture physically supported cross-scale connections, a constrained Local-to-Global interaction is introduced. Only non-center local nodes that satisfy the conditions for motion coupling stability, structural confidence, and dynamics support are allowed to pass information to distal nodes:

$$\widehat{T}_i^G = T_i^G + c_i \,\mathrm{Attn}\left(T_i^G, T_i^L, T_i^L; M_i^{LG}, B_i^{LG}\right), \tag{S7}$$

where $T_i^L$ and $T_i^G$ are the local and distal node representations, respectively, $M_i^{LG}$ is the hard mask for physical availability, and $B_i^{LG}$ is the multiview physical edge bias. The cross-scale gate is defined as

$$c_i = \mathbb{I}\left[\sum_{j,k} M_{i,jk}^{LG} > 0\right] \sigma\left(f_C[h_i^F \| r_i^{LG}]\right), \tag{S8}$$

where $r_i^{LG}$ summarizes the coupling strength, the proportion of valid connections, and structural reliability. When no valid local–distal connections exist, the interaction term is strictly zero.

The base functional representation generates query vectors to retrieve information separately from local nodes and positively and negatively coupled distal nodes. The positive and negative distal retrieval outputs are fused to form global evidence $h_i^G$, and the local retrieval output is denoted by $h_i^L$. The model computes scale weights among three options: no correction, Local, and Global:

$$\begin{aligned} (w_i^0, w_i^L, w_i^G) &= \mathrm{MaskedSoftmax}\left(f_S[h_i^F \| h_i^L \| h_i^G \| q_i]\right), \\ d_i &= w_i^L h_i^L + w_i^G h_i^G, \end{aligned} \tag{S9}$$

where $q_i$ is site-level quality information, and $w_i^0$ corresponds to retaining the base functional representation without applying a dynamics correction.

To constrain adaptation using incremental static structural information, the difference between representations before and after incorporating static three-dimensional information is extracted from the same functional transfer encoder:

$$\delta_i^{3D} = s_i^{\text{post-3D}} - s_i^{\text{pre-3D}}. \tag{S10}$$

Based on the functional representation, the static structural increment, dynamics evidence, and quality information, a conditional adaptation gate is computed as

$$g_i = \sigma\left(f_A[h_i^F \| \delta_i^{3D} \| d_i \| q_i]\right). \tag{S11}$$

The integrated dynamics evidence updates the original functional representation through a conditional low-rank adapter with rank 16:

$$\begin{aligned} \Delta h_i &= m_i(1 - w_i^0) g_i B_D \left[\sigma(W_D d_i + b_D) \odot A_D \,\mathrm{LN}(h_i^F)\right], \\ \widetilde{h}_i^F &= h_i^F + \Delta h_i, \end{aligned} \tag{S12}$$

where $A_D \in \mathbb{R}^{16\times 192}$, $B_D \in \mathbb{R}^{192\times 16}$, $\odot$ denotes element-wise multiplication, and $m_i$ indicates whether a valid dynamics neighborhood exists. The output matrix $B_D$ is initialized to zero, so that the base functional predictions remain unchanged at initialization.

Finally, the frozen functional classification head evaluates the original and updated functional representations separately, and their difference is converted into a bounded correction to the base logit:

$$\Delta z_i = m_i \kappa \tanh\left(\frac{H_F(\widetilde{h}_i^F) - H_F(h_i^F)}{\kappa}\right). \tag{S13}$$

The final prediction logit is $z_i^{\text{final}} = z_i^F + \Delta z_i$, and the corresponding functional probability is $p_i = \sigma(z_i^{\text{final}})$. Here, $\kappa = 0.5$ controls the maximum correction magnitude. When valid dynamics inputs are unavailable, i.e., $m_i = 0$, the model leaves the base prediction unchanged.

#### Model training and evaluation

The model uses a two-stage optimization strategy. In the first stage, the pretrained HieraPTM backbone is frozen, and the LoRA adapters, functional fusion module, and newly constructed classification head are optimized using binary cross-entropy to obtain the base functional predictor. In the second stage, this functional predictor is fixed, and only the multiscale dynamics encoding, cross-scale interaction, and conditional low-rank adaptation modules are trained. The loss function in the dynamics stage combines binary cross-entropy, a logit correction constraint, representation update regularization, and a within-protein site ranking constraint to achieve stable refinement of the base predictions.

For a training batch $\mathcal{B}$, the objective function in the dynamics stage is

$$\begin{aligned} \mathcal{L}_{\text{Dyn}} = \mathcal{L}_{\text{BCE}} &+ \lambda_z \mathcal{L}_z \\ &+ \lambda_h \mathcal{L}_h + \lambda_r \mathcal{L}_{\text{rank}}, \end{aligned} \tag{S14}$$

where $\mathcal{L}_{\text{BCE}}$ is the batch-averaged binary cross-entropy, and $\mathcal{L}_z$ and $\mathcal{L}_h$ constrain the logit correction and functional representation update, respectively:

$$\begin{aligned}\mathcal{L}_z &= \frac{1}{|\mathcal{B}|}\sum_{i\in\mathcal{B}}(\Delta z_i)^2,\\ \mathcal{L}_h &= \frac{1}{192|\mathcal{B}|}\sum_{i\in\mathcal{B}}\|\Delta h_i\|_2^2.\end{aligned} \tag{S15}$$

The regularization coefficients are set to $\lambda_z = 0.05$, $\lambda_h = 0.01$, and $\lambda_r = 0.01$.

The within-protein site ranking constraint is defined as

$$\mathcal{L}_{\text{rank}} = \frac{1}{|\mathcal{P}_{\mathcal{B}}|}\sum_{(i,j)\in\mathcal{P}_{\mathcal{B}}}\text{softplus}\left(z_j^{\text{final}} - z_i^{\text{final}}\right), \tag{S16}$$

where $\mathcal{P}_{\mathcal{B}}$ contains pairs of sites from the same protein within the current batch that satisfy $y_i = 1$ and $y_j = 0$. When no valid site pairs exist, $\mathcal{L}_{\text{rank}}$ is set to zero.

All models are optimized using AdamW, and the best checkpoint is selected based on validation performance. Final results are reported as the mean and standard deviation across nine runs with independently resampled unlabeled backgrounds and separate model training. The standard deviation describes cross-run performance variation under this evaluation protocol. Evaluation metrics include AUROC, AUPRC, Accuracy, Precision, Recall, Specificity, F1-score, and the Matthews correlation coefficient (MCC), with a classification threshold of 0.5 used consistently for threshold-dependent metrics.

### 2 Supplementary Figures

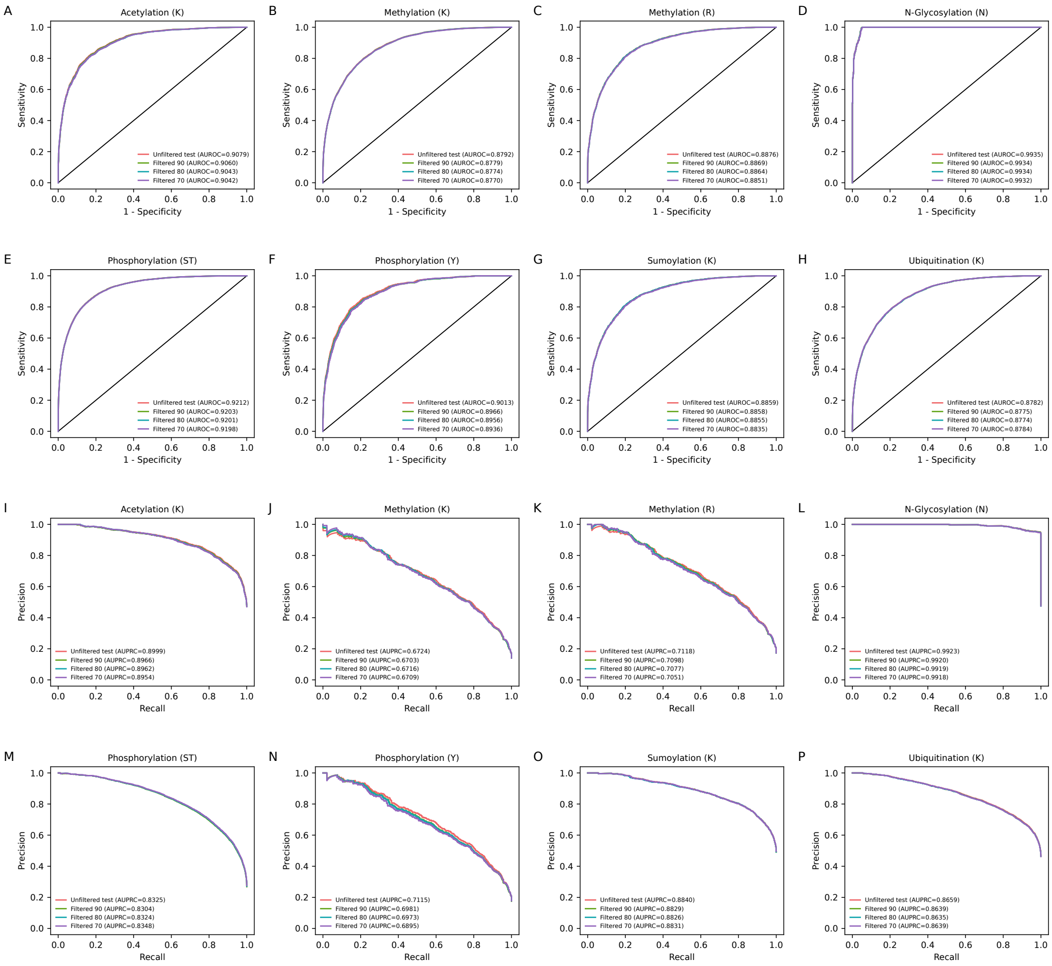


**Fig. S1 Robustness of HieraPTM under protein-disjoint and homology-controlled evaluation.** a–h, ROC curves across eight PTM types on the protein-disjoint test set before and after CD-HIT-2D filtering at 90%, 80%, and 70% sequence identity thresholds. i–p, Corresponding precision–recall curves. AUROC and AUPRC values are shown for each condition.

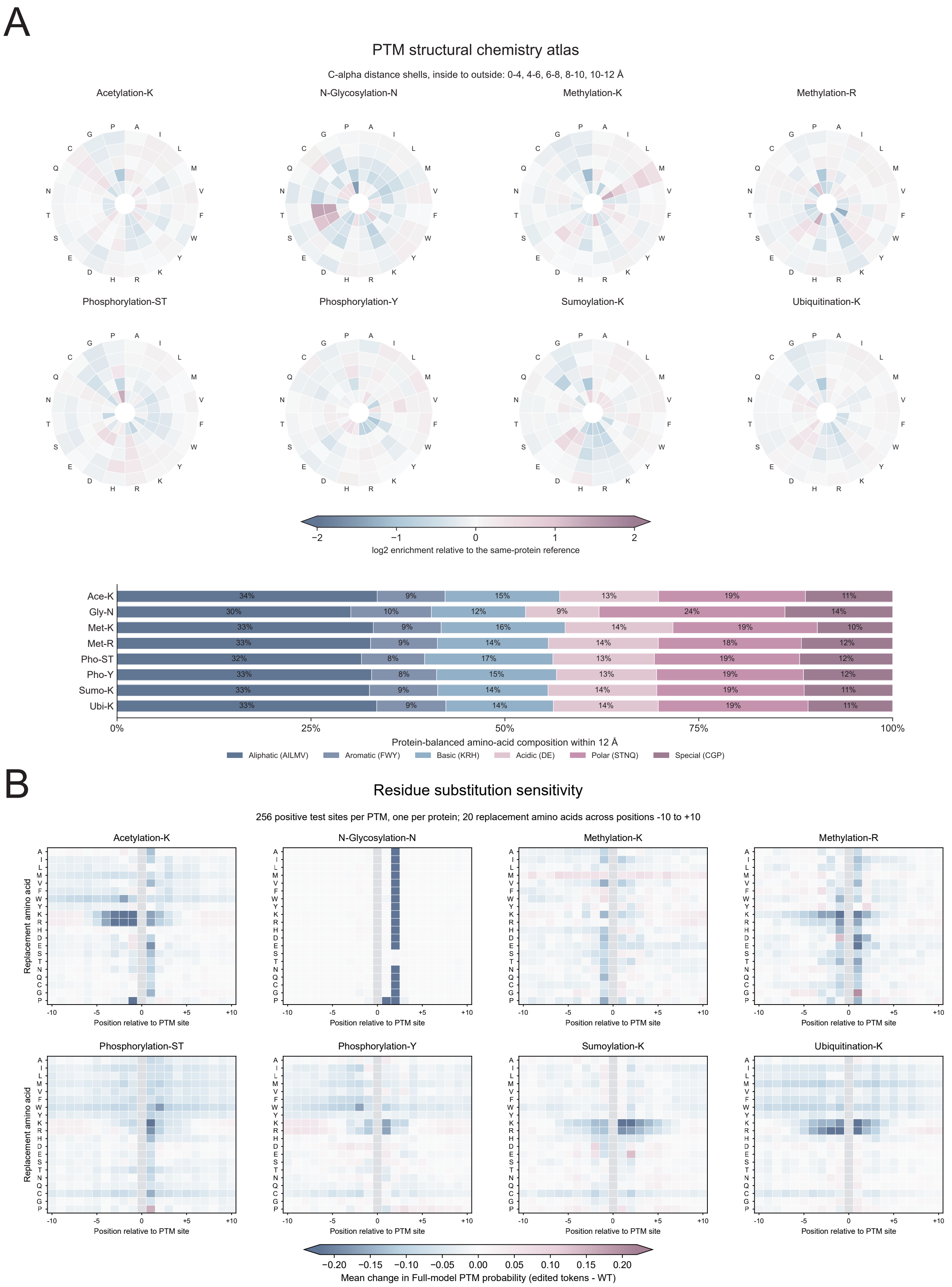


**Fig. S2 PTM-specific structural chemistry and residue substitution sensitivity.** a, Amino acid enrichment across spatial distance shells and biochemical composition of residues within 12 Å of PTM sites across eight PTM types. b, Changes in HieraPTM-predicted PTM probabilities following systematic amino acid substitutions at positions −10 to +10, evaluated on 256 positive test sites per PTM type.

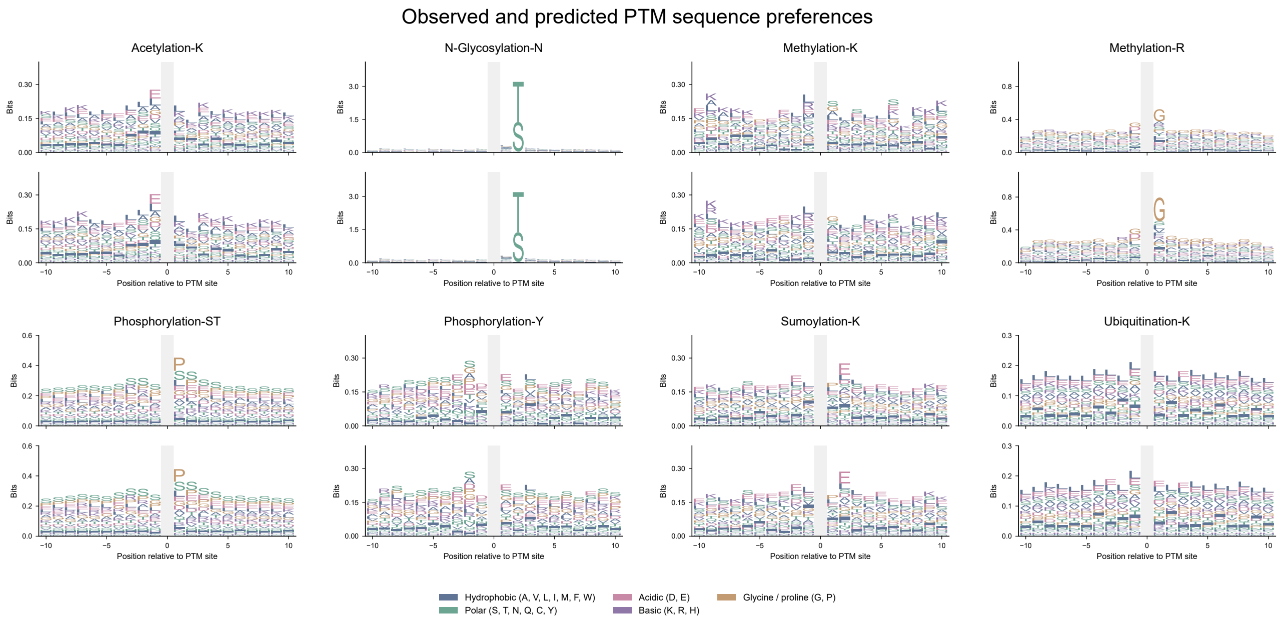


**Fig. S3 Observed and predicted PTM sequence preferences.** Sequence logos of experimentally annotated (top) and HieraPTM-predicted (bottom) sites across eight PTM types.

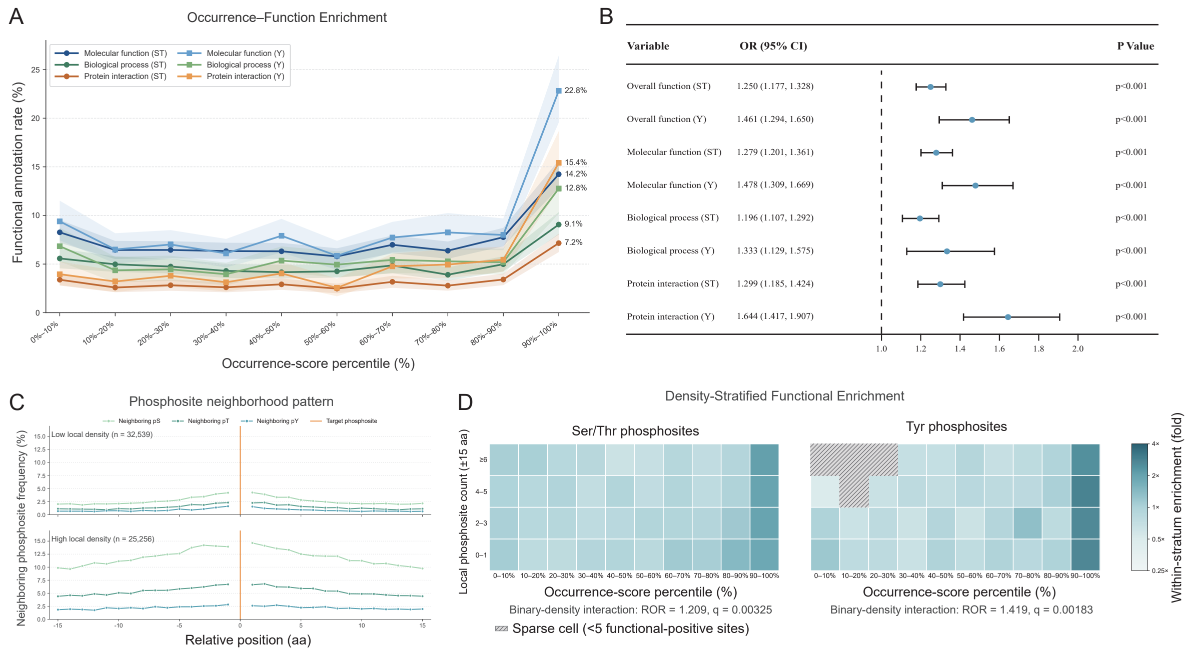


**Fig. S4 HieraPTM site-prediction scores are associated with functional phosphosite annotations.** a, Functional annotation rates across score deciles for three categories in Ser/Thr (ST) and Tyr (Y) phosphosites. Shading indicates 95% protein-bootstrap confidence intervals. b, Within-protein odds ratios (ORs) and 95% confidence intervals for overall and category-specific annotations per standard-deviation increase in prediction score. c, Positional frequencies of neighboring phosphosites within ±15 residues under low-density and high-density conditions. d, Functional annotation enrichment across score deciles and local phosphosite-count strata. Hatched cells indicate fewer than five functionally annotated sites. Interaction ratios of ORs (RORs) and BH-adjusted $q$ values were obtained from binary-density models. All analyses used frozen full HieraPTM scores from 57,795 phosphosites with unseen 61-residue windows, with sequence-only fallback for sites lacking mapped structures.

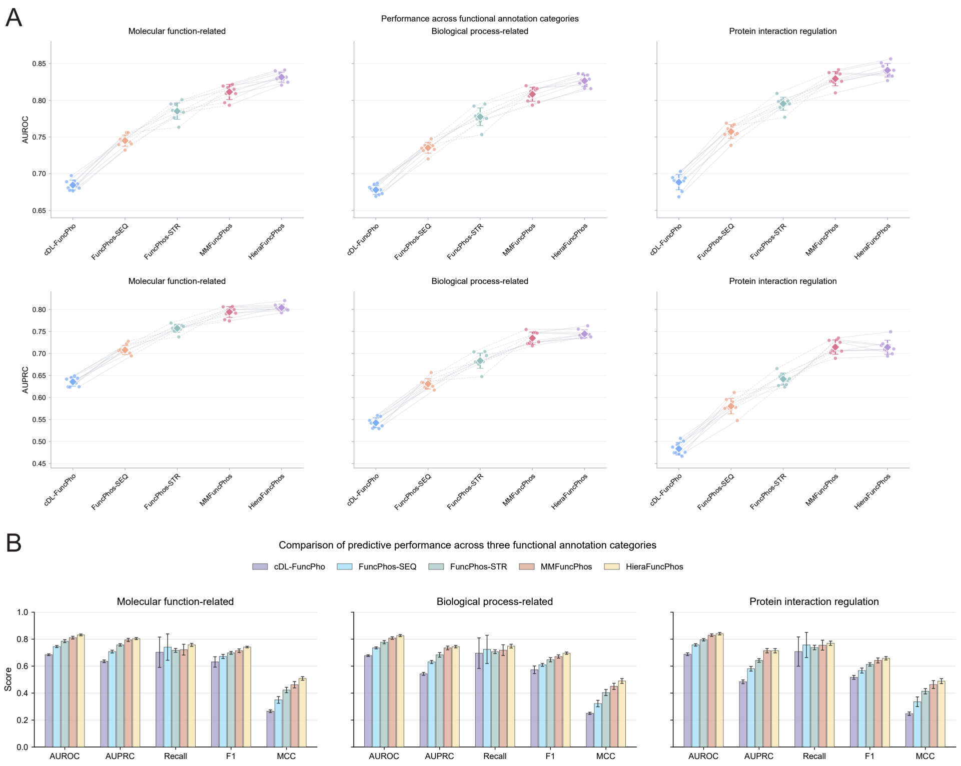


**Fig. S5 Comparative performance of functional phosphosite predictors across three functional annotation categories.** a, AUROC and AUPRC comparisons of five prediction models across molecular function-related, biological process-related, and protein interaction regulation categories. b, Comprehensive comparison of AUROC, AUPRC, Recall, F1-score, and MCC across the three functional annotation categories. The evaluated models include cDL-FuncPhos, FuncPhos-SEQ, FuncPhos-STR, MMFuncPhos, and HieraFuncPhos. All models were evaluated using matched training, validation, and held-out test partitions within each run. Results are summarized across nine runs with independently resampled unlabeled backgrounds and separate model training, with error bars representing cross-run standard deviations.

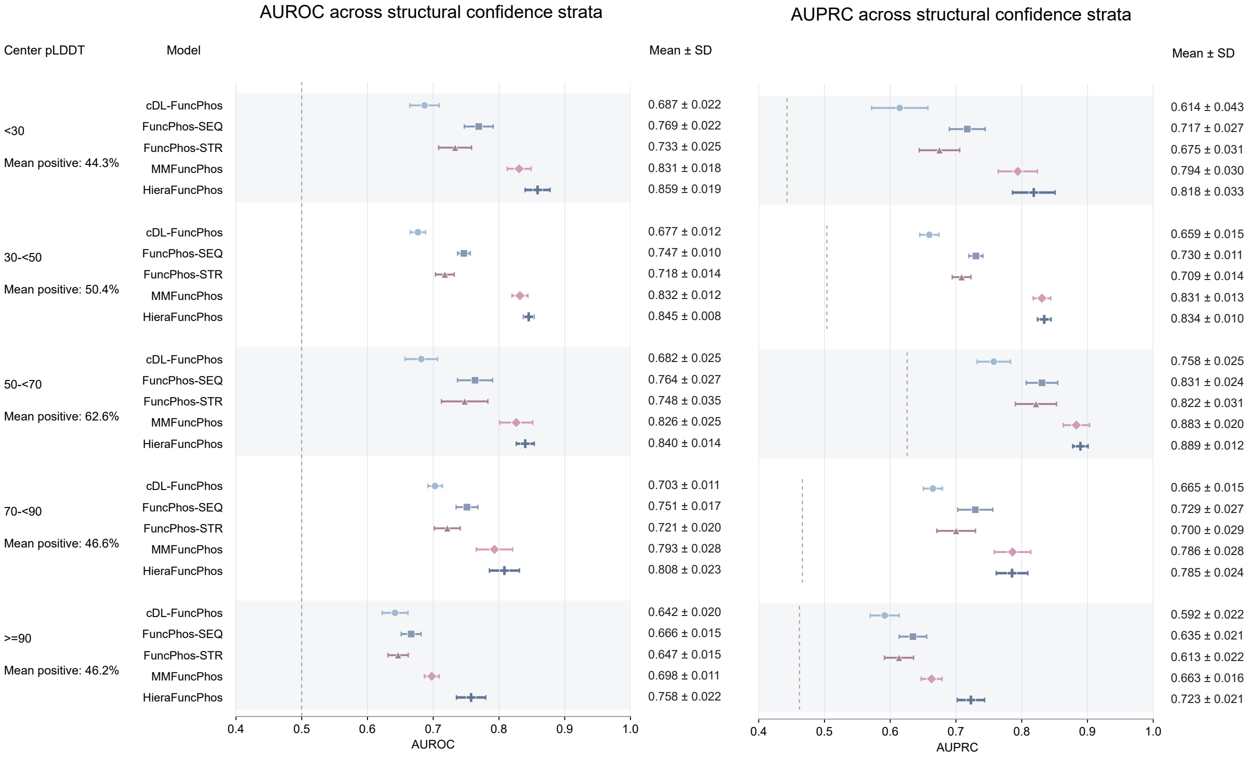


**Fig. S6 Predictive performance across structural confidence strata.** AUROC and AUPRC comparisons of HieraFuncPhos and four baseline models across five center-residue pLDDT intervals. Values represent mean ± standard deviation across repeated experiments.

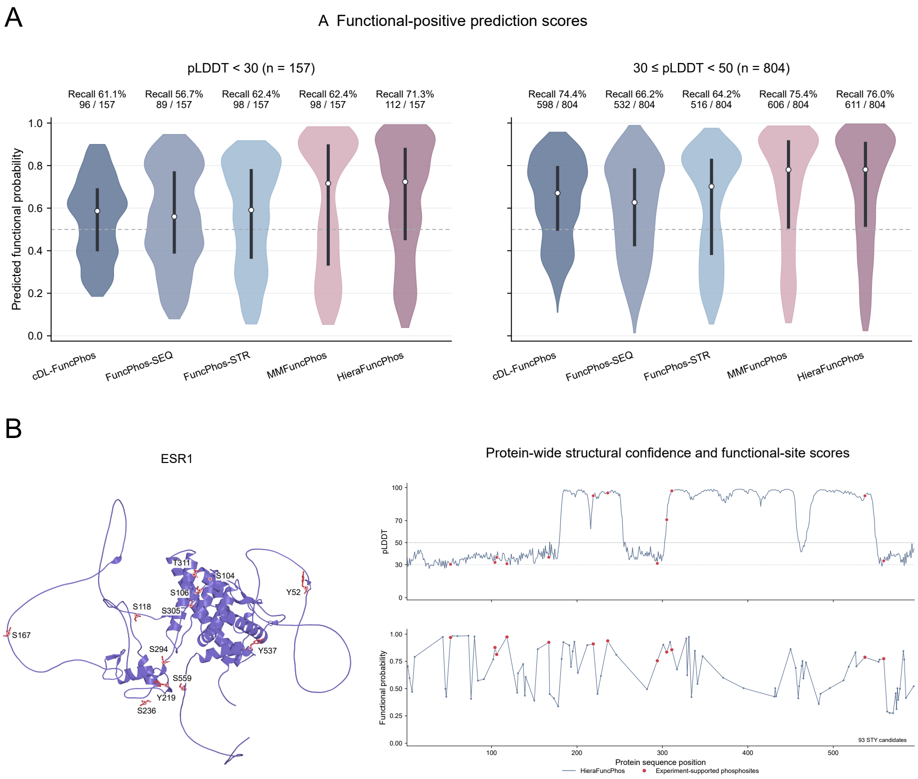


**Fig. S7 Functional phosphosite recognition in regions of low structural confidence.** a, Predicted functional probability distributions across five models for annotated functional phosphosites with pLDDT $< 30$ ($n = 157$) and $30 \leq$ pLDDT $< 50$ ($n = 804$). Recall and correctly identified site counts are reported at a fixed threshold of 0.5. b, Representative ESR1 analysis showing the structural locations of annotated phosphosites, protein-wide pLDDT profiles, and HieraFuncPhos functional predictions across 93 STY candidate sites.

### 3 Supplementary Tables

**Table S1 Summary of the Task 1 PTM-site datasets and multimodal data coverage.**

| PTM task | Total sites | Positive | Putative negative | Proteins | Positive (%) |
|---|---|---|---|---|---|
| N-Glycosylation-N | 14,574 | 7,039 | 7,535 | 2,700 | 48.30 |
| Methylation-K | 39,833 | 5,594 | 34,239 | 2,995 | 14.04 |
| Methylation-R | 49,721 | 8,616 | 41,105 | 3,446 | 17.33 |
| Phosphorylation-Y | 58,765 | 10,396 | 48,369 | 4,390 | 17.69 |
| Acetylation-K | 63,309 | 30,056 | 33,253 | 6,089 | 47.48 |
| Sumoylation-K | 71,468 | 35,248 | 36,220 | 5,725 | 49.32 |
| Ubiquitination-K | 205,487 | 96,132 | 109,355 | 10,512 | 46.78 |
| Phosphorylation-ST | 613,231 | 164,941 | 448,290 | 12,165 | 26.90 |
| **Overall** | **1,116,388** | **358,022** | **758,366** | **16,058**[a] | **32.07** |

[a] The overall protein count represents the number of unique proteins across all eight PTM tasks. Because the same protein may contain sites belonging to multiple PTM tasks, the per-task protein counts are not additive.
All 1,116,388 candidate sites contain a 61-residue local sequence input. Among them, 1,116,190 sites (99.98%) have valid ESM-2 representations and 976,513 sites (87.47%) have available AlphaFold structural information. Samples with unavailable auxiliary modalities were retained and handled using the corresponding modality-validity masks.

**Table S2 Task-specific performance of HieraPTM across eight PTM prediction tasks.**

| PTM task | AUROC | AUPRC | Accuracy (%) | F1 (%) | Precision (%) | Recall (%) | Specificity (%) | MCC |
|---|---|---|---|---|---|---|---|---|
| Acetylation-K | 0.9357 | 0.9274 | 85.61 | 84.79 | 85.12 | 84.46 | 86.65 | 0.7114 |
| N-Glycosylation-N | 0.9969 | 0.9960 | 98.29 | 98.24 | 97.35 | 99.15 | 97.48 | 0.9658 |
| Methylation-K | 0.8908 | 0.6880 | 86.62 | 60.19 | 51.67 | 72.09 | 88.99 | 0.5347 |
| Methylation-R | 0.9032 | 0.7218 | 88.96 | 63.62 | 74.19 | 55.68 | 95.94 | 0.5809 |
| Phosphorylation-ST | 0.9459 | 0.8817 | 88.29 | 79.54 | 75.02 | 84.64 | 89.63 | 0.7163 |
| Phosphorylation-Y | 0.9096 | 0.7293 | 82.00 | 62.13 | 49.49 | 83.46 | 81.68 | 0.5433 |
| Sumoylation-K | 0.9065 | 0.9067 | 83.10 | 82.72 | 83.43 | 82.01 | 84.15 | 0.6619 |
| Ubiquitination-K | 0.9012 | 0.8889 | 81.59 | 81.01 | 78.27 | 83.96 | 79.51 | 0.6334 |

**Table S3 Summary of the dbPTM external-source benchmark under standard and CD-HIT 70% filtering conditions across four PTM prediction tasks.**

| PTM task | Proteins | Positive | Negative | Total sites | Positive (%) | Pos:Neg |
|---|---|---|---|---|---|---|
| *dbPTM* | | | | | | |
| N-Glycosylation-N | 1,018 | 1,332 | 3,904 | 5,236 | 25.44 | 1:2.93 |
| Methylation-K | 2,060 | 3,219 | 14,335 | 17,554 | 18.34 | 1:4.45 |
| Methylation-R | 2,393 | 5,334 | 12,686 | 18,020 | 29.60 | 1:2.38 |
| Ubiquitination-K | 4,265 | 6,106 | 4,265 | 10,371 | 58.88 | 1:0.70 |
| *dbPTM (CD-HIT 70%)* | | | | | | |
| N-Glycosylation-N | 998 | 1,312 | 3,853 | 5,165 | 25.40 | 1:2.94 |
| Methylation-K | 1,959 | 3,023 | 14,084 | 17,107 | 17.67 | 1:4.66 |
| Methylation-R | 2,328 | 5,137 | 12,521 | 17,658 | 29.09 | 1:2.44 |
| Ubiquitination-K | 4,136 | 5,907 | 4,190 | 10,097 | 58.50 | 1:0.71 |

**Table S4 Summary of the PhosF3C-derived external-source benchmark under standard and CD-HIT 70% filtering conditions across two phosphorylation prediction tasks.**

| PTM task | Proteins | Positive | Negative | Total sites | Positive (%) | Pos:Neg |
|---|---|---|---|---|---|---|
| *PhosF3C* | | | | | | |
| Phosphorylation-ST | 461 | 3,671 | 29,566 | 33,237 | 11.04 | 1:8.05 |
| Phosphorylation-Y | 798 | 1,636 | 12,328 | 13,964 | 11.72 | 1:7.54 |
| *PhosF3C (CD-HIT 70%)* | | | | | | |
| Phosphorylation-ST | 451 | 3,613 | 28,957 | 32,570 | 11.09 | 1:8.01 |
| Phosphorylation-Y | 785 | 1,602 | 12,094 | 13,696 | 11.70 | 1:7.55 |

**Table S5: Detailed ablation results of HieraPTM across eight PTM prediction tasks.**

| Configuration | AUROC | AUPRC | Accuracy (%) | F1 (%) | Precision (%) | Recall (%) | Specificity (%) | MCC |
|---|---|---|---|---|---|---|---|---|
| *Acetylation-K* | | | | | | | | |
| **Full** | 0.9356 | 0.9274 | 85.61 | 84.79 | 85.12 | 84.46 | 86.65 | 0.7114 |
| w/o CNN-BiGRU | 0.9207 | 0.9141 | 84.24 | 83.70 | 82.22 | 85.23 | 83.34 | 0.6849 |
| w/o Transformer | 0.9248 | 0.9151 | 84.36 | 84.06 | 81.44 | 86.86 | 82.11 | 0.6888 |
| w/o ESM-2 | 0.9189 | 0.9086[†] | 83.46 | 82.65 | 82.36 | 82.93 | 83.94 | 0.6685 |
| w/o 3D | 0.9313 | 0.9247 | 85.45 | 84.51 | 85.45 | 83.60 | 87.13 | 0.7082 |
| Fixed gate | 0.9231 | 0.9129 | 83.92 | 82.94 | 83.56 | 82.34 | 85.35 | 0.6774 |
| w/o Center Global Access | 0.9310 | 0.9220 | 84.82 | 84.52 | 81.92 | 87.29 | 82.59 | 0.6979 |
| *N-Glycosylation-N* | | | | | | | | |
| **Full** | 0.9969 | 0.9960 | 98.29 | 98.24 | 97.35 | 99.15 | 97.48 | 0.9658 |
| w/o CNN-BiGRU | 0.9952 | 0.9946 | 97.67 | 97.60 | 97.05 | 98.15 | 97.21 | 0.9534 |
| w/o Transformer | 0.9953 | 0.9937 | 98.42 | 98.39 | 97.10 | 99.72 | 97.21 | 0.9688 |
| w/o ESM-2 | 0.9912 | 0.9867 | 98.15 | 98.11 | 96.69 | 99.57 | 96.82 | 0.9634 |
| w/o 3D | 0.9962 | 0.9953 | 98.08 | 98.03 | 96.94 | 99.15 | 97.08 | 0.9618 |
| Fixed gate | 0.9965 | 0.9953 | 98.22 | 98.17 | 97.08 | 99.29 | 97.21 | 0.9646 |
| w/o Center Global Access | 0.9962 | 0.9943 | 98.15 | 98.10 | 97.34 | 98.86 | 97.48 | 0.9630 |
| *Methylation-K* | | | | | | | | |
| **Full** | 0.8908 | 0.6880 | 86.62 | 60.19 | 51.67 | 72.09 | 88.99 | 0.5347 |
| w/o CNN-BiGRU | 0.8738 | 0.6209 | 80.15 | 53.06 | 39.70 | 79.96 | 80.18 | 0.4639 |
| w/o Transformer | 0.8652 | 0.5778 | 86.47 | 56.36 | 51.48 | 62.25 | 90.42 | 0.4874 |
| w/o ESM-2 | 0.8693 | 0.5788 | 88.10 | 58.35 | 57.34 | 59.39 | 92.79 | 0.5142 |
| w/o 3D | 0.8976 | 0.6870 | 90.64 | 62.96 | 70.76 | 56.71 | 96.18 | 0.5814 |
| Fixed gate | 0.8791 | 0.6458 | 85.89 | 58.18 | 49.81 | 69.95 | 88.50 | 0.5103 |
| w/o Center Global Access | 0.8841 | 0.6574 | 80.10 | 53.27 | 39.72 | 80.86 | 79.97 | 0.4677 |
| *Methylation-R* | | | | | | | | |
| **Full** | 0.9032 | 0.7218 | 88.96 | 63.62 | 74.19 | 55.68 | 95.94 | 0.5809 |
| w/o CNN-BiGRU | 0.8749 | 0.6533 | 81.94 | 59.37 | 48.66 | 76.10 | 83.17 | 0.5047 |
| w/o Transformer | 0.8802 | 0.6751 | 85.12 | 63.04 | 55.35 | 73.20 | 87.62 | 0.5477 |
| w/o ESM-2 | 0.8798 | 0.6560 | 85.26 | 62.47 | 55.91 | 70.77 | 88.30 | 0.5403 |
| w/o 3D | 0.8907 | 0.6884 | 87.73 | 62.67 | 66.32 | 59.40 | 93.68 | 0.5548 |
| Fixed gate | 0.8911 | 0.6889 | 88.34 | 57.29 | 78.43 | 45.13 | 97.40 | 0.5372 |
| w/o Center Global Access | 0.8959 | 0.7020 | 87.95 | 65.11 | 65.38 | 64.85 | 92.80 | 0.5784 |
| *Phosphorylation-ST* | | | | | | | | |
| **Full** | 0.9459 | 0.8817 | 88.29 | 79.54 | 75.02 | 84.64 | 89.63 | 0.7163 |
| w/o CNN-BiGRU | 0.9371 | 0.8622 | 86.39 | 77.24 | 70.20 | 85.86 | 86.59 | 0.6838 |
| w/o Transformer | 0.9305 | 0.8502 | 86.24 | 76.57 | 70.63 | 83.61 | 87.21 | 0.6741 |
| w/o ESM-2 | 0.9317 | 0.8556 | 86.70 | 76.97 | 72.02 | 82.66 | 88.18 | 0.6800 |
| w/o 3D | 0.9460 | 0.8814 | 88.13 | 79.39 | 74.50 | 84.96 | 89.30 | 0.7141 |
| Fixed gate | 0.9356 | 0.8602 | 86.62 | 77.14 | 71.35 | 83.95 | 87.60 | 0.6821 |
| w/o Center Global Access | 0.9438 | 0.8760 | 86.69 | 78.04 | 70.16 | 87.92 | 86.24 | 0.6957 |

*Continued on next page*

*Continued from previous page*

| Configuration | AUROC | AUPRC | Accuracy (%) | F1 (%) | Precision (%) | Recall (%) | Specificity (%) | MCC |
|---|---|---|---|---|---|---|---|---|
| *Phosphorylation-Y* | | | | | | | | |
| **Full** | 0.9096 | 0.7293 | 82.00 | 62.13 | 49.49 | 83.46 | 81.68 | 0.5433 |
| w/o CNN-BiGRU | 0.8905 | 0.6814 | 81.93 | 60.67 | 49.34 | 78.75 | 82.61 | 0.5202 |
| w/o Transformer | 0.8968 | 0.6977 | 82.18 | 61.63 | 49.79 | 80.87 | 82.47 | 0.5341 |
| w/o ESM-2 | 0.8883 | 0.6770 | 82.47 | 58.73 | 50.34 | 70.48 | 85.05 | 0.4909 |
| w/o 3D | 0.9046 | 0.7167 | 81.49 | 61.34 | 48.65 | 82.98 | 81.17 | 0.5333 |
| Fixed gate | 0.9049 | 0.7164 | 80.60 | 60.58 | 47.30 | 84.23 | 79.82 | 0.5262 |
| w/o Center Global Access | 0.9049 | 0.7129 | 78.00 | 58.65 | 43.94 | 88.17 | 75.81 | 0.5103 |
| *Sumoylation-K* | | | | | | | | |
| **Full** | 0.9065 | 0.9067 | 83.10 | 82.72 | 83.43 | 82.01 | 84.15 | 0.6619 |
| w/o CNN-BiGRU | 0.8897 | 0.8858 | 80.87 | 80.86 | 79.84 | 81.90 | 79.87 | 0.6177 |
| w/o Transformer | 0.8884 | 0.8898 | 80.48 | 80.59 | 79.08 | 82.16 | 78.85 | 0.6102 |
| w/o ESM-2 | 0.8783 | 0.8717 | 79.50 | 79.43 | 78.63 | 80.26 | 78.77 | 0.5902 |
| w/o 3D | 0.9063 | 0.9064 | 82.66 | 82.19 | 83.30 | 81.11 | 84.18 | 0.6533 |
| Fixed gate | 0.8938 | 0.8938 | 81.25 | 80.42 | 82.92 | 78.07 | 84.35 | 0.6257 |
| w/o Center Global Access | 0.9039 | 0.9034 | 82.76 | 82.83 | 81.42 | 84.28 | 81.28 | 0.6557 |
| *Ubiquitination-K* | | | | | | | | |
| **Full** | 0.9012 | 0.8889 | 81.59 | 81.01 | 78.27 | 83.96 | 79.51 | 0.6334 |
| w/o CNN-BiGRU | 0.8881 | 0.8741 | 80.15 | 79.34 | 77.31 | 81.47 | 78.99 | 0.6034 |
| w/o Transformer | 0.8890 | 0.8713 | 80.47 | 79.70 | 77.56 | 81.97 | 79.15 | 0.6100 |
| w/o ESM-2 | 0.8867 | 0.8658 | 79.71 | 78.58 | 77.62 | 79.57 | 79.83 | 0.5932 |
| w/o 3D | 0.9008 | 0.8884 | 81.33 | 81.15 | 76.90 | 85.89 | 77.32 | 0.6315 |
| Fixed gate | 0.8919 | 0.8755 | 80.69 | 79.89 | 77.89 | 82.00 | 79.54 | 0.6142 |
| w/o Center Global Access | 0.9000 | 0.8876 | 81.11 | 80.96 | 76.59 | 85.85 | 76.94 | 0.6274 |

**Table S6 Run-wise predictive performance of HieraFuncPhos under repeated background sampling and model retraining.**

| Rep | AUROC | AUPRC | ACC (%) | Precision (%) | Recall (%) | Specificity (%) | F1 (%) | MCC |
|---|---|---|---|---|---|---|---|---|
| 1 | 0.8348 | 0.8225 | 75.44 | 76.52 | 73.42 | 77.47 | 74.94 | 0.5093 |
| 2 | 0.8264 | 0.8184 | 75.22 | 74.55 | 76.58 | 73.86 | 75.55 | 0.5046 |
| 3 | 0.8287 | 0.8166 | 75.14 | 74.28 | 76.91 | 73.36 | 75.57 | 0.5031 |
| 4 | 0.8330 | 0.8206 | 75.42 | 76.17 | 73.97 | 76.86 | 75.06 | 0.5085 |
| 5 | 0.8165 | 0.8073 | 73.78 | 73.10 | 75.25 | 72.31 | 74.16 | 0.4758 |
| 6 | 0.8240 | 0.8201 | 75.03 | 76.10 | 72.97 | 77.08 | 74.50 | 0.5010 |
| 7 | 0.8374 | 0.8328 | 75.86 | 76.06 | 75.47 | 76.25 | 75.77 | 0.5172 |
| 8 | 0.8257 | 0.8159 | 75.25 | 75.28 | 75.19 | 75.31 | 75.24 | 0.5050 |
| 9 | 0.8206 | 0.8151 | 74.25 | 74.22 | 74.31 | 74.20 | 74.27 | 0.4850 |

**Table S7 Ablation study configuration of hierarchical knowledge transfer, functional adaptation, and mechanical refinement.**

| Model | Residue ESM-2 | Static structure graph | Protein-global ESM-2 | PTM transfer | Functional LoRA | Mechanical retrieval |
|---|---|---|---|---|---|---|
| Model 1 | × | × | × | × | × | × |
| Model 2 | ✓ | × | × | × | × | × |
| Model 3 | ✓ | ✓ | ✓ | × | × | × |
| Model 4 | ✓ | ✓ | ✓ | ✓ | × | × |
| Model 5 | ✓ | ✓ | ✓ | ✓ | ✓ | × |
| **Model 6** | ✓ | ✓ | ✓ | ✓ | ✓ | ✓ |

**Table S8 Ablation analysis of hierarchical knowledge transfer, functional adaptation, and mechanical refinement.**

| Model | AUROC | AUPRC | ACC (%) | Recall (%) | Specificity (%) | F1 (%) | MCC |
|---|---|---|---|---|---|---|---|
| Model 1 | 0.7059±0.0118 | 0.6893±0.0146 | 65.04±1.12 | 62.10±5.47 | 67.98±6.18 | 63.90±2.03 | 0.3032±0.0228 |
| Model 2 | 0.7482±0.0053 | 0.7323±0.0046 | 68.38±0.38 | 70.77±4.94 | 65.98±4.99 | 69.05±1.65 | 0.3695±0.0071 |
| Model 3 | 0.7826±0.0067 | 0.7648±0.0078 | 71.34±0.59 | 70.07±4.50 | 72.61±3.46 | 70.91±1.70 | 0.4280±0.0116 |
| Model 4 | 0.8174±0.0068 | 0.8092±0.0083 | 74.15±0.73 | 74.03±2.43 | 74.26±2.91 | 74.11±0.81 | 0.4835±0.0144 |
| Model 5 | 0.8225±0.0071 | 0.8149±0.0071 | 74.43±0.74 | 74.07±1.70 | 74.79±2.49 | 74.34±0.62 | 0.4890±0.0149 |
| **Model 6** | **0.8274±0.0068** | **0.8188±0.0068** | **75.04±0.64** | **74.90±1.35** | **75.19±1.84** | **75.01±0.59** | **0.5011±0.0128** |

**Table S9 Mechanical refinement gains across baseline prediction confidence quintiles.**

| Baseline Confidence Quintile | ΔAUROC (95% CI) | ΔAUPRC (95% CI) |
|---|---|---|
| Q1 (Highest confidence) | +0.0020 [0.0003, 0.0038] | +0.0002 [−0.0017, 0.0021] |
| Q2 (High confidence) | +0.0073 [0.0037, 0.0110] | +0.0054 [−0.0012, 0.0126] |
| Q3 (Intermediate confidence) | +0.0188 [0.0136, 0.0240] | +0.0211 [0.0119, 0.0297] |
| Q4 (Low confidence) | +0.0280 [0.0219, 0.0338] | +0.0308 [0.0208, 0.0398] |
| Q5 (Lowest confidence) | +0.0358 [0.0281, 0.0436] | +0.0390 [0.0286, 0.0479] |

**Table S10 Training hyperparameters and computational costs of HieraPTM and HieraFuncPhos across different training stages.**

| Hyperparameter | Sequence pretraining | Structural fusion | Functional LoRA | Dynamic adaptation |
|---|---|---|---|---|
| Optimizer | AdamW | Grouped AdamW | Grouped AdamW | AdamW |
| Batch size | 64 | 64 | 32 | 32 |
| Maximum epochs | 80 | 30 | 60 | 40 |
| Early-stopping patience | 10 | 6 | 10 | 8 |
| Weight decay | $1 \times 10^{-4}$ | $1 \times 10^{-4}$ | $1 \times 10^{-2}$ | $1 \times 10^{-2}$ |
| Gradient clipping | 1 | 1 | 1 | 1 |
| Training loss | Weighted BCE | Weighted BCE + residual regularization | BCE | BCE + dynamic regularization |
| *Learning-rate configuration* | | | | |
| Base learning rate | $3 \times 10^{-4}$ | Group-specific | Group-specific | $3 \times 10^{-4}$ |
| Graph encoder | – | $5 \times 10^{-5}$ | – | – |
| Cross-attention | – | $1 \times 10^{-4}$ | – | – |
| Gates, output projection and normalization | – | $2 \times 10^{-4}$ | – | – |
| LoRA parameters | – | – | $1 \times 10^{-4}$ | – |
| ESM fusion gate and structural residual coefficients | – | – | $3 \times 10^{-5}$ | – |
| Global projection, functional adapter and classifier | – | – | $3 \times 10^{-4}$ | – |
| *Computational cost* | | | | |
| Trainable parameters | 2,719,970 | | 1,639,900 | |
| Cumulative training time | 10.89 h | | 1.87 h | |
| Peak training GPU memory | 0.474 GB | | 0.141 GB | |

*Note:* Sequence pretraining and structural fusion correspond to the two training stages of HieraPTM, while functional LoRA and dynamic adaptation correspond to the two training stages of HieraFuncPhos. LR denotes learning rate; BCE denotes binary cross-entropy; LoRA denotes low-rank adaptation. A dash indicates that the corresponding parameter group is not applicable. Cumulative training time represents the total training duration across the corresponding stages, excluding feature extraction and cache construction. All experiments were conducted on an NVIDIA GeForce RTX 4090 GPU. Peak training GPU memory denotes the maximum GPU memory consumption recorded during model training.
